# A bacterial effector protein prevents MAPK-mediated phosphorylation of SGT1 to suppress plant immunity

**DOI:** 10.1101/641241

**Authors:** Gang Yu, Liu Xian, Hao Xue, Wenjia Yu, Jose S. Rufian, Yuying Sang, Rafael J. L. Morcillo, Yaru Wang, Alberto P. Macho

## Abstract

Nucleotide-binding domain and leucine-rich repeat-containing (NLR) proteins function as sensors that perceive pathogen molecules and activate immunity. In plants, the accumulation and activation of NLRs is regulated by SUPPRESSOR OF G2 ALLELE OF *skp1* (SGT1). In this work, we found that an effector protein named RipAC, secreted by the plant pathogen *Ralstonia solanacearum*, associates with SGT1 to suppress NLR-mediated SGT1-dependent immune responses, including those triggered by another *R. solanacearum* effector, RipE1. RipAC does not affect the accumulation of SGT1 or NLRs, or their interaction. However, RipAC inhibits the interaction between SGT1 and MAP kinases, and the phosphorylation of a MAPK target motif in the C-terminal domain of SGT1. Such phosphorylation is enhanced upon activation of immune signaling, leads to the release of the interaction between SGT1 and NLRs, and contributes to the activation of NLR-mediated responses. Additionally, SGT1 phosphorylation contributes to resistance against *R. solanacearum*, and this is particularly evident in the absence of RipAC. Our results shed light onto the mechanism of activation of NLR-mediated immunity, and suggest a positive feedback loop between MAPK activation and SGT1-dependent NLR activation.

## Introduction

The activation and suppression of plant immunity are key events that determine the outcome of the interaction between plants and bacterial pathogens. When in close proximity to plant cells, conserved bacterial molecules (termed PAMPs, for pathogen-associated molecular patterns) may be perceived by receptors at the surface of plant cells, eliciting the activation of the plant immune system. This leads to the establishment of PAMP-triggered immunity (PTI; Boller and Felix, 2009), which restricts bacterial growth and prevents the development of disease. Most bacterial pathogens have acquired the ability to deliver proteins inside plant cells via a type-III secretion system (T3SS); such proteins are thus called type-III effectors (T3Es) (Galán et al., 2014). Numerous T3Es from adapted bacterial plant pathogens have been reported to suppress PTI (Macho and Zipfel, 2015). This, in addition to other T3E activities (Macho, 2016), enables bacterial pathogens to proliferate and cause disease. Resistant plants have evolved mechanisms to detect such bacterial manipulation through the development of intracellular receptors defined by the presence of nucleotide-binding sites (NBS) and leucine-rich repeat domains (LRRs), thus termed NLRs (Cui et al., 2015). The detection of T3E activities through NLRs leads to the activation of immune responses, which effectively prevent pathogen proliferation (Chiang and Coaker, 2015). The outcome of these responses is named effector-triggered immunity (ETI), and, in certain cases, may cause a hypersensitive response (HR) that involves the collapse of plant cells. In an evolutionary response to this phenomenon, T3E activities have evolved to suppress ETI (Jones and Dangl, 2006), which in turn exposes bacterial pathogens to further events of effector recognition. For these reasons, the interaction between plants and microbial pathogens is often considered an evolutionary ‘arms race’, where the specific combination of virulence activities and immune receptors in a certain pathogen-plant pair ultimately defines the outcome of the interaction (Jones and Dangl, 2006). Although the suppression of PTI by T3Es has been widely documented (Macho and Zipfel, 2015), reports about T3Es that suppress ETI, and their biochemical characterization, remain scarce.

Interestingly, PTI and ETI signaling often involve shared regulators (Tsuda and Katagiri, 2010), such as mitogen-activated protein kinases (MAPK). The activation of MAPKs is one of the earliest signaling events upon perception of both PAMPs and effectors (Meng and Zhang, 2013), and is essential for the activation of immune responses. Although sustained MAPK activation has been reported to be critical for the robust development of ETI (Tsuda et al., 2013), the signaling events and phosphorylation substrates that link early MAPK activation upon pathogen recognition and downstream MAPK activation as a consequence of NLR activation remain unknown.

The study of microbial effectors has become a very prolific approach to discover and characterize components of the plant immune system and other cellular functions that play important roles in plant-microbe interactions (Toruño et al., 2016). In this regard, T3E repertoires constitute a powerful tool for researchers to understand plant-bacteria interactions. The *Ralstonia solanacearum* species complex (RSSC) groups numerous bacterial strains able to cause disease in more than 250 plant species, including important crops, such as tomato, potato, and pepper, among others (Jiang et al., 2017; Mansfield et al., 2012). *R. solanacearum* invades the roots of host plants, reaching the vascular system and colonizing xylem vessels in the whole plant, eventually causing their blockage and subsequent plant wilting (Digonnet et al., 2012; Turner et al., 2009). It has been speculated that the versatility and wide host range of *R. solanacearum* is associated to its relatively large T3E repertoire in comparison with other bacterial plant pathogens; the reference GMI1000 strain serves as an example, being able to secrete more than 70 T3Es (Peeters et al., 2013). Among these T3Es, RipAC (also called PopC; Guéneron et al., 2000; Peeters et al., 2013) is conserved in most *R. solanacearum* sequenced to date, and is therefore considered a core T3E in the RSSC (Figures S1A and S1B) (Peeters et al., 2013). Domain prediction for the RipAC amino acid sequence reveals the presence of an LRR C-terminal domain and no other predicted enzymatic domain (Figure S1C). Interestingly, it has been shown that RipAC from different *R. solanacearum* strains is promptly expressed upon bacterial inoculation in different host plants (Jacobs et al., 2012; Puigvert et al., 2017) suggesting a role in the establishment of the interaction at early stages of the infection process.

## Results

### RipAC contributes to *Ralstonia solanacearum* infection

It is well established that T3Es collectively play an essential role in the development of disease caused by most bacterial plant pathogens, including *R. solanacearum* (Deslandes and Genin, 2014). To study the contribution of RipAC to *R. solanacearum* infection, we generated a *ΔripAC* mutant strain using the reference phylotype I GMI1000 strain as background (Figure S1D), and confirmed that the growth of this strain in rich laboratory medium is indistinguishable from the wild-type (WT) strain (Figures S1D-1G). We then performed soil-drenching inoculation of Arabidopsis Col-0 WT plants with both WT and *ΔripAC* strains, and monitored the development of wilting symptoms associated to disease. Compared with plants inoculated with WT GMI1000, plants inoculated with the *ΔripAC* mutant displayed attenuated disease symptoms (Figures 1A, S2A-2C). This attenuation was complemented by the expression of *ripAC*, driven by its native promoter, in the *ΔripAC* mutant background (Figures 1A, S2A-2C, and S1E-1G), confirming that virulence attenuation was indeed caused by the lack of the *ripAC* gene. Given the intrinsic complexity and biological variability of the soil-drenching inoculation assay, we did not detect a significant attenuation every time we performed this experiment (Figure S2C), suggesting that the attenuation caused by the absence of RipAC is near the limit of detection of this assay. Tomato is a natural host for several *R. solanacearum* strains (Hayward, 1991). As observed in Arabidopsis, tomato plants inoculated with the *ΔripAC* strain showed delayed and/or reduced symptom development in most experiments (Figures 1B, S2A, S2B and S2D). This attenuation was complemented by the expression of *ripAC* in the mutant background (Figures 1B, S2A, S2B and S2D).

**Figure 1.**
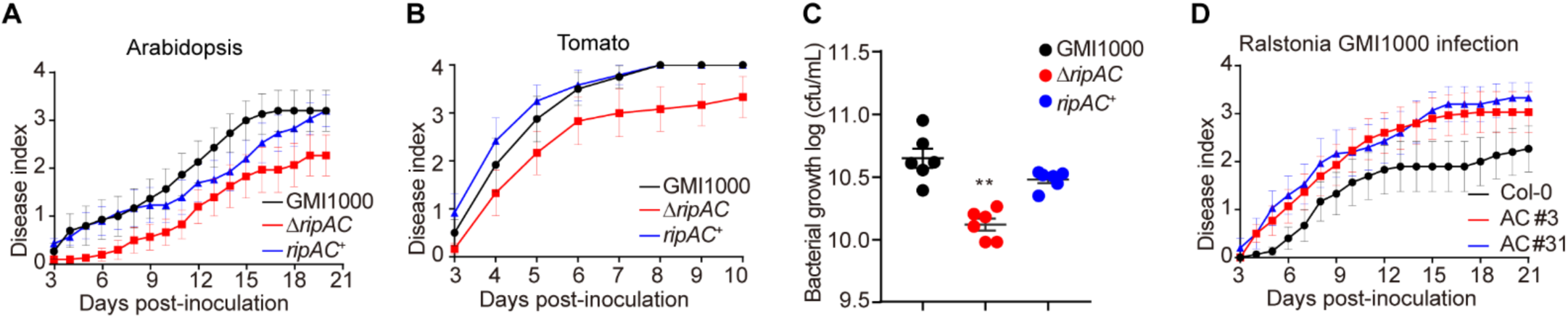
RipAC contributes to *Ralstonia solanacearum* virulence in Arabidopsis and tomato. (A, B) Soil-drenching inoculation assays in Arabidopsis (A) and tomato (B) were performed with GMI1000 WT, Δ*ripAC* mutant, and RipAC complementation (*ripAC*^+^) strains. n=15 plants per genotype (for Arabidopsis) or n=12 plants per genotype (for tomato). The results are represented as disease progression, showing the average wilting symptoms in a scale from 0 to 4 (mean ± SEM). (C) Growth of *R. solanacearum* GMI1000 WT, Δ*ripAC* mutant, and RipAC complementation (*ripAC*^+^) strains in tomato plants. 5 µL of bacterial inoculum (10^6^ cfu mL^−1^) were injected into the stem of 4-week-old tomato plants and xylem sap was taken from each infected plants for bacterial titer calculation 3 days post-inoculation (dpi) (mean ± SEM, n=6, ** p<0.01, *t*-test). (D) Soil-drenching inoculation assays in Col-0 WT and RipAC-expressing Arabidopsis lines (AC #3, AC #31, two independent lines) with GMI1000 WT strain. These experiments were repeated at least 3 times with similar results. See figures S2 and S3 for independent replicates and a composite representation.

In order to increase the accuracy of our virulence assays and to determine whether the reduced disease symptoms correlate with reduced bacterial multiplication inside host plants, we set up an assay to quantify bacterial numbers in the xylem sap of tomato plants upon injection of the bacterial inoculum in the stem. Compared to plants inoculated with WT GMI1000 or the complementation strain, plants inoculated with the *ΔripAC* strain showed reduced bacterial numbers (Figure 1C). Altogether, these results suggest that, despite the large effector repertoire of *R. solanacearum* GMI1000, RipAC plays a significant role during the infection process in different host plants.

To complement our genetic analysis of the importance of RipAC in promoting susceptibility to *R. solanacearum*, we generated stable Arabidopsis transgenic lines expressing RipAC from a 35S constitutive promoter (Figures S3A and S3B). Independent transgenic lines expressing RipAC showed enhanced disease symptoms upon soil-drenching inoculation with *R. solanacearum* (Figures 1D and S3C-D), indicating that RipAC promotes susceptibility to *R. solanacearum* from within plant cells.

### RipAC interacts with SGT1 from different plant species

Upon transient expression in *Nicotiana benthamiana*, RipAC fused to a C-terminal GFP tag shows a sharp signal at the cell periphery of leaf epidermal cells (Figure S4A). Fractionation of protein extracts from *N. benthamiana* expressing RipAC, showed abundant RipAC in both microsomal and cytosolic fractions (Figure S4B). To identify plant proteins that physically associate with RipAC in plant cells, we performed immunoprecipitation (IP) of RipAC-GFP using GFP-trap agarose beads, and analyzed its interacting proteins using liquid chromatography coupled to tandem mass-spectrometry (LC-MS/MS). Together with RipAC-GFP immunoprecipitates, we identified peptides of NbSGT1 (Figure S5A), homolog of the Arabidopsis SUPPRESSOR OF G2 ALLELE OF *skp1* (SGT1), a protein required for the induction of disease resistance mediated by most NLRs (Azevedo et al., 2002; Kadota et al., 2010). The localization of both isoforms of SGT1 in Arabidopsis (AtSGT1a and AtSGT1b) has been reported to be nucleocytoplasmic (Noël et al., 2007). When we expressed SGT1 (NbSGT1, AtSGT1a, or AtSGT1b) fused to a C-terminal RFP tag, we also detected an apparently cytoplasmic localization, with occasional nuclear localization, although we also detected co-localization with the plasma membrane marker CBL-GFP (Gao et al., 2011; Figures S5B and S5C). Interestingly, we detected co-localization of RipAC-GFP and NbSGT1-RFP, AtSGT1a-RFP, or AtSGT1b-RFP (Figure S5D). These results also suggest that RipAC does not affect SGT1 subcellular localization. To confirm the physical interaction between RipAC and SGT1 from different plant species, we co-expressed RipAC-GFP with either NbSGT1, AtSGT1a, AtSGT1b or the tomato ortholog of SGT1b (SlSGT1b). Targeted IP of RipAC-GFP showed CoIP of all different SGT1 orthologs upon transient co-expression in *N. benthamiana* (Figure 2A). Similarly, split-luciferase complementation (Split-LUC) assays in *N. benthamiana* showed luciferase signal upon expression of RipAC with AtSGT1a, AtSGT1b or NbSGT1 fused to different luciferase halves (Figure 2B). We also performed split-YFP assays, showing reconstitution of the YFP signal at the cell periphery upon expression of RipAC and AtSGT1a, AtSGT1b or NbSGT1 tagged with one of the portions of a split-YFP molecule (Figure S5E), indicating a direct interaction of these proteins at the cell periphery. Altogether, these results indicate that RipAC physically associates with SGT1 from different species inside plant cells.

**Figure 2.**
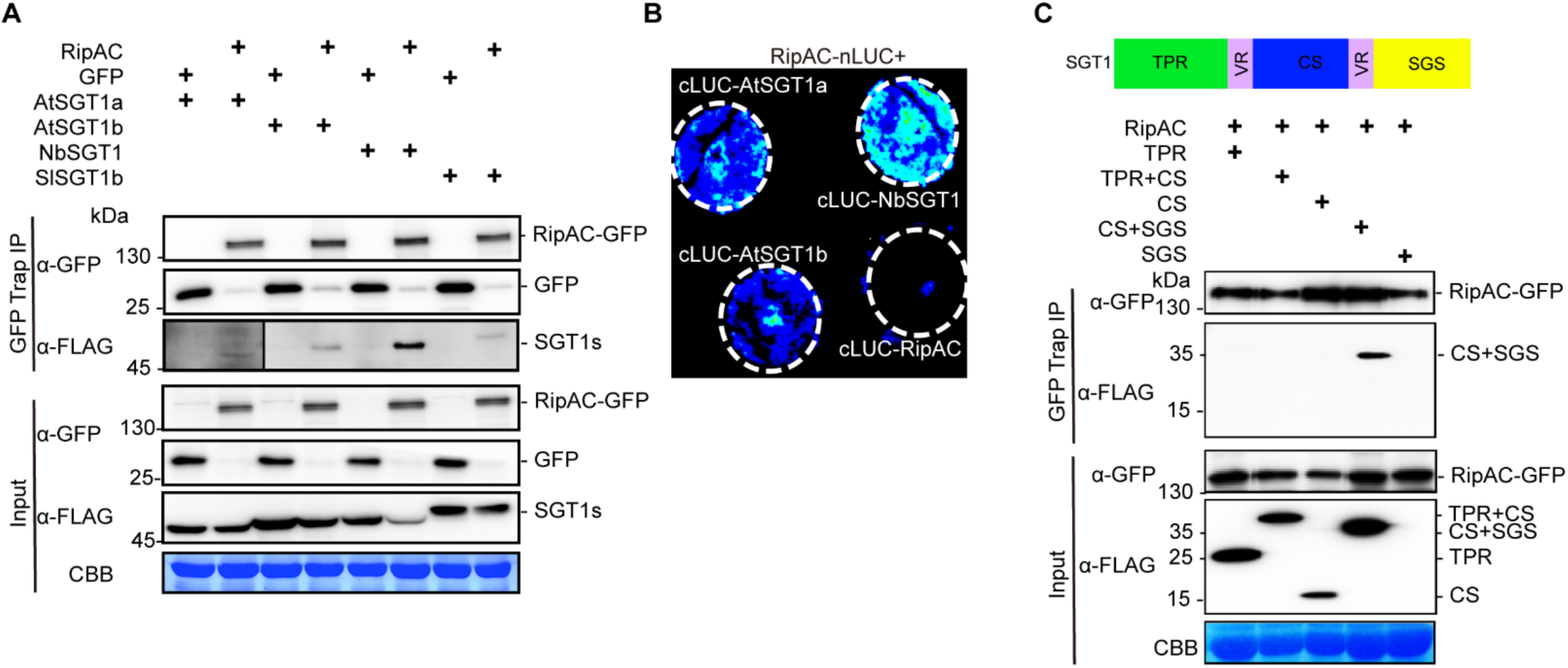
RipAC associates with SGT1 in plant cells. (A) CoIP to determine interactions between RipAC and SGT1s. The signal in the interaction between RipAC and AtSGT1a was weak and is shown with longer exposure. (B) Split-LUC assay to determine direct interaction between RipAC and SGT1 in plant cells. (C) CoIP to determine interactions between RipAC and different truncated versions of NbSGT1. The diagram summarizes the different domains of SGT1: TPR, N-terminal tetratricopeptide repeat (TPR) domain; CS, CHORD-SGT1 domain; SGS, C-terminal SGT1-specific domain; VR, variable region. Experiments in (A) and (B) were repeated at least 3 times with similar results. The experiment in (C) was repeated 2 times with similar results.

SGT1 is conserved in eukaryotes, and has three defined domains: an N-terminal tetratricopeptide repeat (TPR) domain, a CHORD-SGT1 (CS) domain, and a C-terminal SGT1-specific (SGS) domain (Shirasu, 2009). In order to determine which NbSGT1 domain is required for RipAC interaction, we expressed different combinations of these domains (Figure 2C) bound to a FLAG tag, and tested their interaction with immunoprecipitated RipAC-GFP. Although we were unable to express the SGS domain alone, we detected interaction between RipAC and a truncated SGT1 protein containing the CS+SGS domain (Figure 2C). The CS domain alone did not interact with RipAC, suggesting that RipAC associates with the SGS domain, although it cannot be ruled out that RipAC requires both CS+SGS domains to associate with SGT1. Interestingly, the SGS domain is required for all the known SGT1b functions in immunity (Noël et al., 2007; Shirasu, 2009).

SGT1 physically associates with several NLRs (Bieri et al., 2004; Kud et al., 2013; Leister et al., 2005); here we found that AtSGT1b also interacts with the NLR RPS2 (Bent et al., 1994) (Figure S5F). To determine whether RipAC interferes with the interaction between SGT1 and NLRs, we performed Split-LUC assays between AtSGT1b and RPS2 in the presence of RipAC. Interestingly, RipAC did not affect RPS2 accumulation or the interaction between AtSGT1b and RPS2 (Figures S5F and S5G).

### RipAC inhibits SGT1-mediated immune responses in *N. benthamiana* and Arabidopsis

To determine whether RipAC has the potential to suppress SGT1-mediated immune responses, we first performed transient expression of RipAC-GFP in *N. benthamiana*, and monitored the induction of cell death associated to the hypersensitive response (HR) developed as part of an ETI response, such as that triggered by overexpression of the NLR-encoding gene *RPS2* (Jin et al., 2002), the expression of the oomycete effector Avr3a together with the cognate NLR R3a (Armstrong et al., 2005), and that triggered by inoculation with *P. syringae* pv. *tomato* DC3000 (*Pto* DC3000) (Wei et al., 2007). RipAC suppressed the HR triggered by RPS2 and Avr3a/R3a, while it attenuated and delayed HR development in response to *Pto* DC3000 (Figure 3A), showing the potential of RipAC to suppress ETI. We have recently reported that the *R. solanacearum* T3E RipE1 triggers SGT1-dependent immunity in *N. benthamiana* (Sang et al., 2019). RipAC was also able to suppress RipE1-induced HR (Figure 3A). Interestingly, RipAC was not able to suppress the cell death triggered by other elicitors, such as Bax (Lacomme and Santa Cruz, 1999) and the oomycete elicitor INF1 (Kamoun et al., 1998) (Figure 3B), suggesting that RipAC is not a general suppressor of cell death. Ion leakage measurements validated that RipAC suppresses SGT1-dependent ETI-associated cell death (e.g. triggered by RPS2 or RipE1), but had a minor impact on BAX-triggered cell death (Figures S6A and S6B).

**Figure 3.**
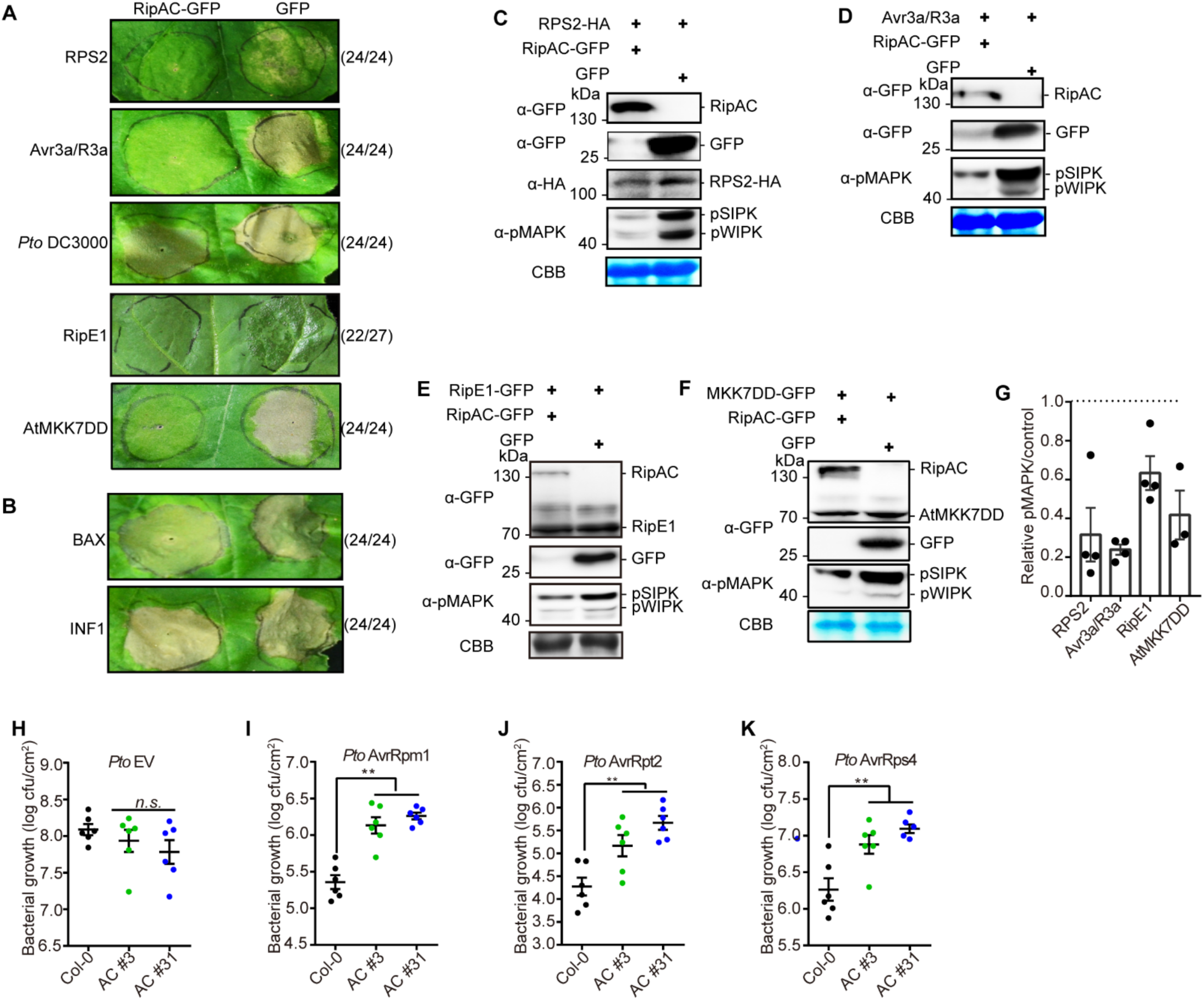
RipAC inhibits SGT1-dependent immune responses. (A) RipAC suppresses RPS2-, Avr3a/R3a-, *Pto* DC3000-, RipE1-, or AtMKK7DD-associated cell death. (B) RipAC does not suppress BAX- or INF1-induced cell death. Photographs were taken 5 days after the expression of the cell death inducer. The numbers beside the photographs indicate the ratios of infiltration showing the presented result among the total number of infiltrations. (C-G) RipAC suppresses MAPK activation triggered by the overexpression of RPS2 (C), Avr3a/R3a (D), RipE1 (E) or AtMKK7DD (F) in *Nicotiana benthamiana*. Plant tissues were taken one day after cell death inducer expression. (G) Quantification of relative pMAPK in (C-F). The quantification of the pMAPK band in RipAC expression spots is represented relative to the intensity in the GFP control in each assay (mean ± SEM of 3 or 4 independent biological replicates). (H-K) RipAC suppresses effector-triggered immunity in Arabidopsis. Leaves of Col-0 WT and RipAC-transgenic lines (AC #3 and AC #31) were hand-infiltrated with *Pto* empty vector (EV, H), *Pto* AvrRpm1 (I), *Pto* AvrRpt2 (J), or *Pto* AvrRps4 (K). Samples were taken 3dpi (mean ± SEM, n=6, ** p<0.01, *t*-test). These experiments were repeated at least 3 times with similar results.

The activation of NLRs is known to trigger the activation of mitogen-activated protein kinases (MAPKs; Adachi et al., 2015; Tsuda et al., 2013). Consistently, transient overexpression of RPS2 in *N. benthamiana* leads to an activation of MAPKs (SIPK and WIPK) that precedes the onset of ion leakage and macroscopic cell death (Figures S6A-S6D). In our transient expression assays, RipAC significantly suppressed the activation of MAPKs triggered by overexpression of RPS2, Avr3a/R3a, or by the expression of RipE1 (Figures 3C-E and 3G), supporting the notion that RipAC suppresses early steps in ETI activation rather than downstream responses that lead to cell death. In agreement with this, RipAC also abolished the macroscopic cell death and MAPK activation triggered by the overexpression of a constitutively active form of the Arabidopsis MAPK Kinase 7 (AtMKK7DD) in *N. benthamiana* (Figures 3A, 3F, and 3G), which requires SGT1 function (Popescu et al., 2009). These results indicate that RipAC is able to suppress ETI-associated SGT1-dependent responses from different origins in *N. benthamiana*.

To determine whether RipAC inhibits the final outcome of ETI (i.e. restriction of bacterial proliferation), we inoculated RipAC-expressing Arabidopsis transgenic lines (Figures S3A and S3B) with *Pto* DC3000 expressing different T3Es that activate NLR-dependent immunity, namely AvrRpm1 (Grant et al., 1995), AvrRpt2 (Bent et al., 1994), or AvrRps4 (Gassmann et al., 1999). Such strains, compared to *Pto* DC3000 expressing an empty vector (EV) (Figure 3H), show deficient replication in Arabidopsis, due to immune responses triggered by these T3Es (Figures 3I-3K). However, transgenic lines expressing RipAC showed enhanced susceptibility to these strains (Figures 3I-3K), indicating that RipAC is able to effectively suppress NLR-mediated immunity in Arabidopsis.

### SGT1 is phosphorylated in a MAPK target motif upon activation of ETI

Since post-translational modifications (PTMs) have been shown to impact SGT1 function (Hoser et al., 2013; Kim et al., 2014; Redkar et al., 2015), we sought to determine whether RipAC causes any alteration of SGT1 PTMs in plant cells. To this end, we co-expressed NbSGT1 fused to a FLAG tag together with RipAC (or a GFP control) in *N. benthamiana* leaves, and then performed IP using an anti-FLAG resin followed by LC-MS/MS. Among the IPed SGT1 peptides, we detected two peptides, located in the SGS domain, containing phosphorylated serine residues (S282 and S358) (Figure S7). Interestingly, a phosphorylable residue is conserved in these positions among plants from different species (Figure S8A), and S358 is embedded in a canonical MAPK-mediated phosphorylation motif conserved across the plant kingdom (Figure S8B). Interestingly, AtSGT1b has been shown to interact with MAPK4 (Cui et al., 2010), and SGT1 from tobacco (NtSGT1) S358, equivalent to T346 in AtSGT1b, has been shown to be phosphorylated by NtSIPK (Hoser et al., 2013). By performing targeted CoIP, we found that both AtSGT1a and AtSGT1b associate with AtMAPK4 and AtMAPK6, but not with AtMAPK3, which showed very low accumulation in this assay (Figure 4A). In Split-LUC assays, we found direct interaction of both AtSGT1a and AtSGT1b with AtMAPK3 and AtMAPK4, but not with AtMAPK6 (Figures 4B, 4C, and S9). These results indicate that AtMAPK4 interacts with AtSGT1a and AtSGT1b. While they also suggest an interaction of AtSGT1s with AtMAPK3 and AtMAPK6, different results were obtained in different interaction assays, probably due to the different experimental conditions. To determine whether MAPK-mediated SGT1 phosphorylation is enhanced upon activation of ETI, we raised an antibody against the potential MAPK phosphorylated peptide, conserved in both AtSGT1a and AtSGT1b, containing phosphorylated T346 (ESpTPPDGME), which specifically detected the phosphorylated peptide *in vitro* and *in planta* (Figures S10A and S10B). Inoculation of Arabidopsis plants with *Pto* expressing AvrRpt2 or AvrRps4, but not with *Pto* DC3000 or *Pto* TTSS^−^, led to a sustained activation of MAPKs and to enhanced phosphorylation of T346 (Figure S10C). Similarly, dexamethasone-induced expression of AvrRpt2 in Arabidopsis transgenic lines led to sustained MAPK activation and enhanced SGT1 phosphorylation (Figures 4D and 4E). These data indicate that an enhancement of T346 phosphorylation occurs concomitantly to the activation of ETI.

**Figure 4.**
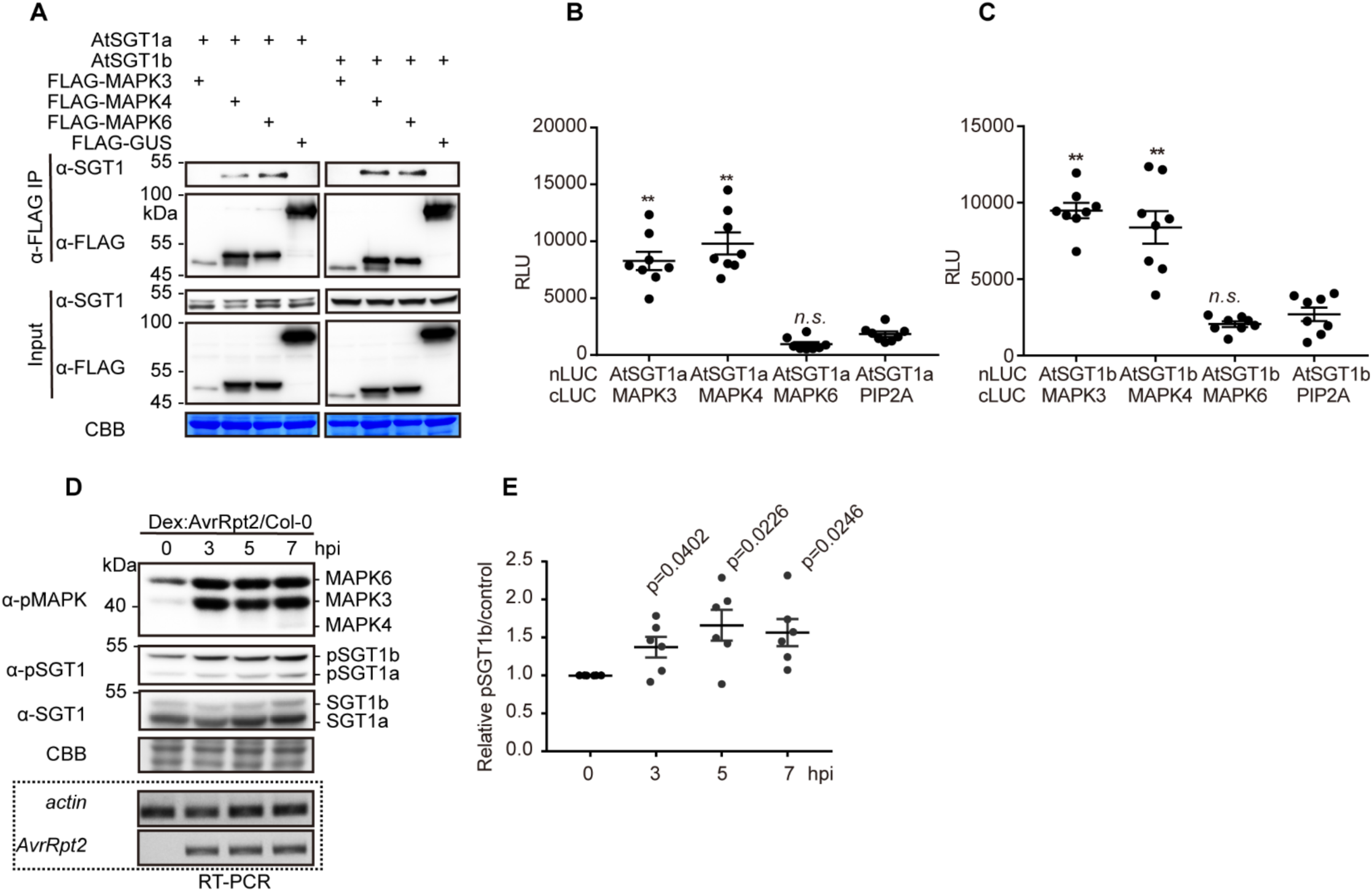
MAPKs associate with SGTs in plant cells. (A) AtSGT1a and AtSGT1b associate with MAPK4 and MAPK6 in plant cells. CoIPs were performed as in Fig. 2A. (B,C) AtSGT1a/b associate with AtMAPK3/4, but not with MAPK6, in Split-LUC assay. Agrobacterium combinations with different constructs were infiltrated in *N. benthamiana* leaves and luciferase activities were examined with microplate luminescence reader. The graphs show the mean value ± SEM (n=8, ** p<0.01, *t*-test). The MAPK-PIP2A combination was used as negative control. (D) MAPK activation triggered by AvrRpt2 expression correlates with increased AtSGT1 phosphorylation at T346. Twelve-day-old Dex:AvrRpt2/Col-0 crossing F1 plants were treated with 30 μM Dex, and samples were taken at the indicated time points. In (D), a custom antibody was used to detect the phosphorylated peptide containing AtSGT1 T346. The induction of AvrRpt2 was determined by RT-PCR. (E) Quantification of pSGT1b signal relative to coomassie blue staining control in (D) (mean ± SEM of 6 independent biological repeats, p-values are shown, *t*-test). These experiments were repeated at least 3 times with similar results.

### MAPK-mediated phosphorylation is important for SGT1 function in the activation of ETI

To determine the relevance of the phosphorylation of SGT1 S271/T346 for the activation of ETI, we generated single and double point mutants in these residues in AtSGT1b, substituting them for alanine residues (to abolish phosphorylation) or for aspartate residues (to mimic the negative charge associated to constitutive phosphorylation). Due to the difficulties in performing genetic analysis of SGT1, we used a transient approach in *N. benthamiana*, co-expressing the AtSGT1b variants together with RPS2. Alanine mutations (S271A/T346A: AtSGT1b AA) did not have a detectable impact on the RPS2-mediated cell death (Figure 5A), likely due to the presence of endogenous NbSGT1. However, tissues expressing both the T346D mutant (2D) or the double S271D/T346D (DD) mutant, but not the single S271D mutant, showed an enhancement of cell death triggered by RPS2 expression, monitored as ion leakage (Figure 5A), which correlated with the appearance of tissue necrosis (Figures 5B and 5C). Similarly, co-expression with the 2D mutant enhanced cell death triggered by RipE1 and Avr3a/R3a (Figures S10D and S10E), but not BAX (Figure S10F). This suggests that T346 phosphorylation contributes to the robust activation of SGT1-dependent ETI responses, including that triggered by the *R. solanacearum* T3E RipE1.

**Figure 5.**
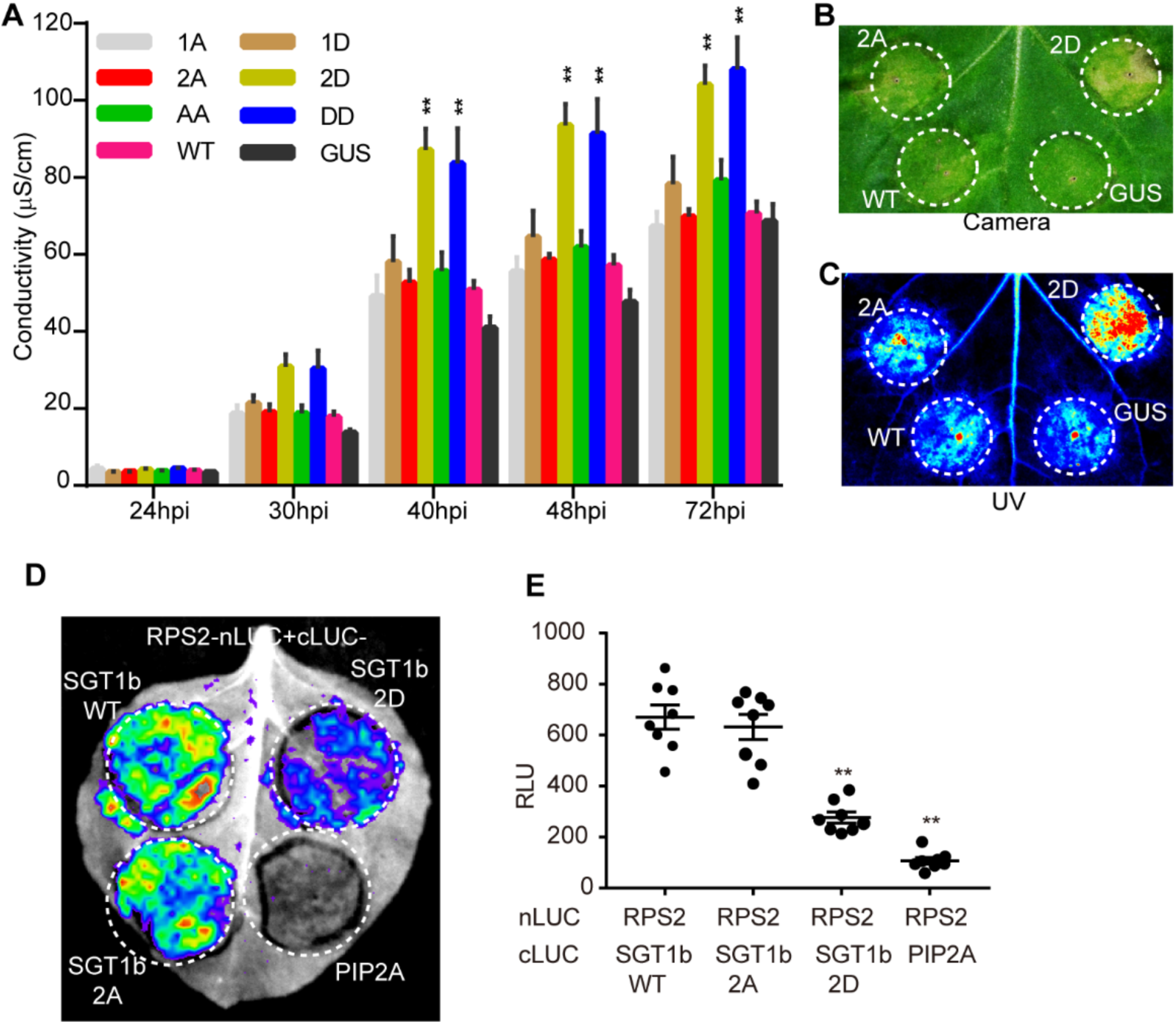
MAPK-mediated phosphorylation is important for SGT1 function in the activation of ETI. (A) A phospho-mimic mutation in AtSGT1b T346 (T346D) promotes cell death triggered by RPS2 overexpression. Agrobacterium expressing AtSGT1b variants or the GUS-FLAG control (OD600=0.5) were infiltrated into *N. benthamiana* leaves 1 day before infiltration with Agrobacterium expressing RPS2 (OD600=0.15). Leaf discs were taken 21 hpi for conductivity measurements at the indicated time points. The time points in the x-axis are indicated as hpi with Agrobacterium expressing RPS2 (mean ± SEM, n=4, ** p<0.01, *t*-test, 3 replicates). (B, C) Observation of RPS2-triggered cell death by visible light (B) and UV light (C). The Agrobacterium combinations were infiltrated as described in (A), and the cell death phenotype was recorded 4 dpi with Agrobacterium expressing RPS2. (D, E) A phospho-mimic mutation in AtSGT1b T346 (T346D) disrupts RPS2-SGT1b association. Agrobacterium combinations with different constructs were infiltrated in *N. benthamiana* leaves and luciferase activities were examined in both qualitative (CCD imaging machine, D) and quantitative assays (microplate luminescence reader, E). Quantification of the luciferase signal was performed with microplate luminescence reader (mean ± SEM, n=8, ** p<0.01, *t*-test, 3 replicates). The nomenclature of the AtSGT1b mutants used is: 1A=S271A, 1D=S271D, 2A=T346A, 2D=T346D, AA=S271AT346A, DD=S271DT346D. The experiments were performed at least 3 times with similar results.

Given the importance of protein-protein interactions in the SGT1 complex, we hypothesized that phosphorylation in S271/T346 may be important to regulate the interaction of SGT1 with NLRs. Interestingly, the T346A mutation did not affect the interaction between AtSGT1b and RPS2 (Figures 5D, 5E, and S10G); however, surprisingly, the T346D mutation inhibited this interaction (Figures 5D, 5E, and S10G). This, together with the observation that the T346D mutation causes an enhancement of RPS2-triggered responses, suggests a scenario where phosphorylation contributes to the release of RPS2 from the SGT1 complex to activate immune responses.

### RipAC interferes with the MAPK-SGT1 interaction to suppress SGT1 phosphorylation

During our analysis of SGT1 phosphorylation, we observed an apparent reduction in the number of SGT1 phosphorylated peptides in the presence of RipAC (Figure S7A). Since this assay does not allow a comparative quantitative assessment of SGT1 phosphorylation, we took advantage of the sensitivity of our anti-pSGT1 antibody to determine the impact of RipAC on the phosphorylation of AtSGT1 T346. Interestingly, Arabidopsis transgenic lines expressing RipAC showed reduced phosphorylation of T346 in endogenous SGT1 in basal growth conditions (Figures 6A and 6B). Such reduction of T346 phosphorylation was also observed when Arabidopsis protoplasts were transfected with RipAC in the presence of AvrRpt2 (Figures S11A and S11B). We then generated Arabidopsis transgenic lines expressing RipAC and dexamethasone-inducible AvrRpt2 by crossing. RipAC completely abolished MAPK activation and T346 phosphorylation induced by the expression of AvrRpt2 (Figures 6C and 6D). Altogether, these data indicate that RipAC suppresses the phosphorylation of AtSGT1 T346, and provides an additional link between MAPK activation and SGT1 phosphorylation. We considered the possibility that RipAC suppresses SGT1 phosphorylation by interfering with the MAPK-SGT1 interaction. This interference was indeed confirmed by Split-LUC (for MAPK3 and MAPK4) and CoIP assays (for MAPK6), which showed that RipAC inhibits the interaction of SGT1 with MAPKs (Figures 6E-6G, S11C-S11F), suggesting that this could be the mechanism underlying RipAC suppression of SGT1-dependent ETI responses (Figure S12).

**Figure 6.**
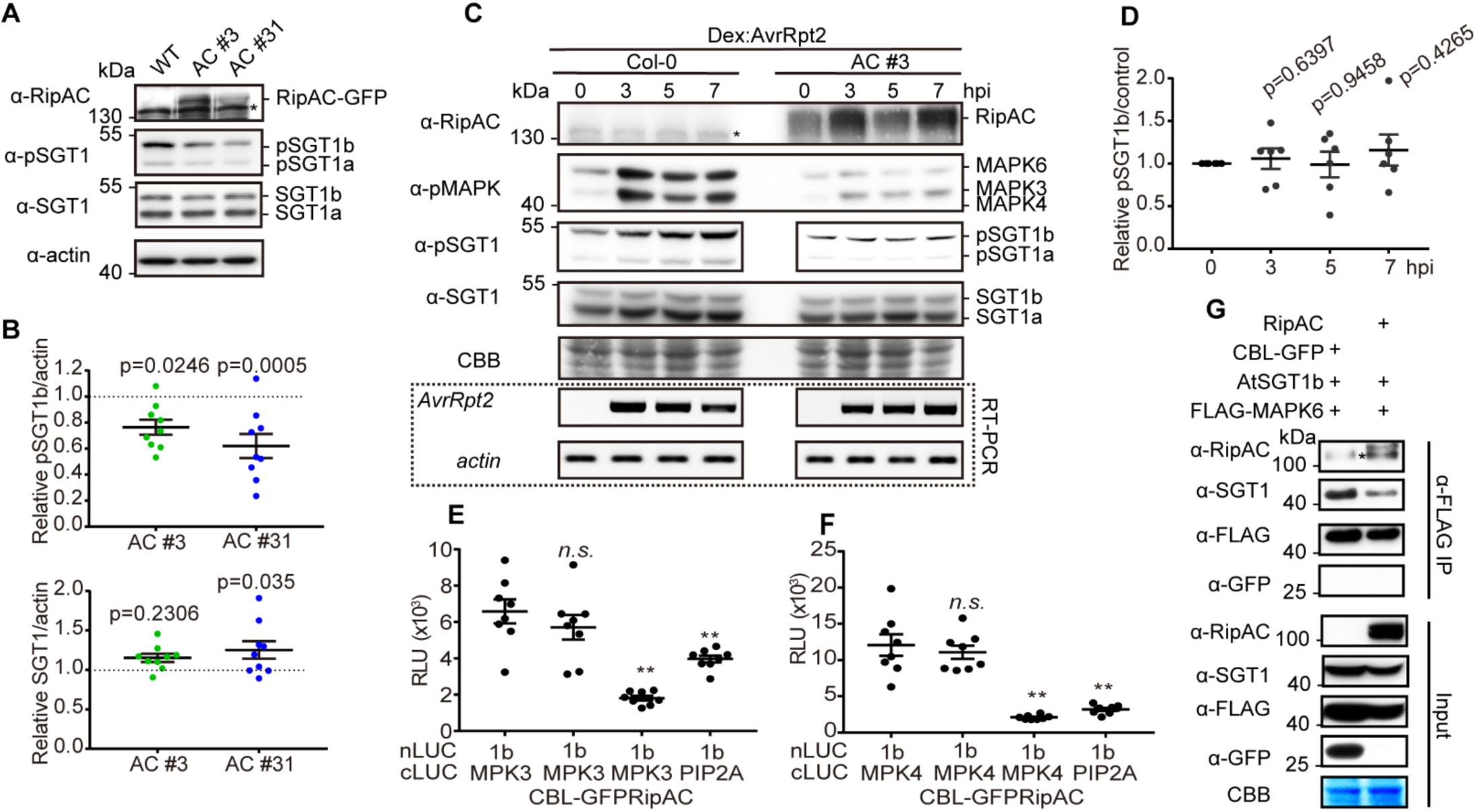
RipAC interferes with MAPK-SGT1 interaction to suppress SGT1 phosphorylation. (A) RipAC reduces SGT1 phosphorylation in Arabidopsis. SGT1 phosphorylation was determined in twelve-day-old Col-0 WT and RipAC transgenic lines (AC #3 and AC #31) by western blot. Custom antibodies were used to detect the accumulation of SGT1 and the presence of a phosphorylated peptide containing AtSGT1 T346. (B) Quantification of SGT1 and pSGT1b signal normalized to actin and relative to Col-0 WT samples in (A) (mean ± SEM, n=9, p-values are shown, *t*-test). (C) RipAC suppresses ETI-triggered SGT1 phosphorylation in Arabidopsis. ETI triggered MAPK activation and SGT1 phosphorylation were determined in twelve-day-old Dex::*AvrRpt2*/Col-0 or Dex::*AvrRpt2*/RipAC crossing F1 plants by western blot. Custom antibodies were used to detect the accumulation of SGT1 and the presence of a phosphorylated peptide containing AtSGT1 T346. The induction of AvrRpt2 was determined by RT-PCR. (D) Quantification of pSGT1b signal relative to coomassie blue staining control in (C) (mean ± SEM of 6 independent biological replicates, p-values are shown, *t*-test). (E, F) Competitive Split-LUC assays showing that RipAC interferes with the interaction between MAPK3 (E) / MAPK4 (F) and AtSGT1b. In quantitative assays, the graphs show the mean value ± SEM (n=8, ** p<0.01, *t*-test). (G) Competitive CoIP showing that RipAC interferes with the interaction between MAPK6 and AtSGT1b. Anti-FLAG beads were used to IP MAPK6. In all the competitive interaction assays, in addition to the interaction pair, RipAC or CBL-GFP (as negative control) were expressed to determine interference. In all the blots, asterisks indicate non-specific bands. These experiments were repeated at least 3 times with similar results.

### SGT1 phosphorylation contributes to resistance against *R. solanacearum* in Arabidopsis

Finally, we sought to determine the biological relevance of SGT1 phosphorylation for RipAC virulence activity during *R. solanacearum* infection. Given the impossibility to isolate *Atsgt1a/b* double mutants, we generated Arabidopsis transgenic lines expressing, from a *35S* promoter, AtSGT1b WT, AtSGT1b 2A, or AtSGT1b 2D, abolishing or mimicking phosphorylation in T346, respectively. Overexpression of AtSGT1b WT did not significantly affect disease symptoms caused by *R. solanacearum* GMI1000 WT (Figures 7A, S13A, and S13B); however, it significantly enhanced resistance against the *ΔripAC* mutant (Figures 7B, S13C, and S13D). Overexpression of AtSGT1b 2A enhanced plant susceptibility against *R. solanacearum* GMI1000 WT (Figures 7A, S13A, and S13B), suggesting that it may exert a dominant negative effect over the endogenous SGT1, and partially rescued the virulence attenuation of the *ΔripAC* mutant (Figures 7B, S13C, and S13D), indicating that SGT1 phosphorylation is relevant for the attenuation phenotype in the absence of RipAC. Intriguingly, overexpression of AtSGT1b 2D slightly enhanced resistance against *R. solanacearum* GMI1000 WT (Figures 7A, S13A, and S13B), and dramatically enhanced resistance against the *ΔripAC* mutant (Figures 7B, S13C, and S13D).

**Figure 7.**
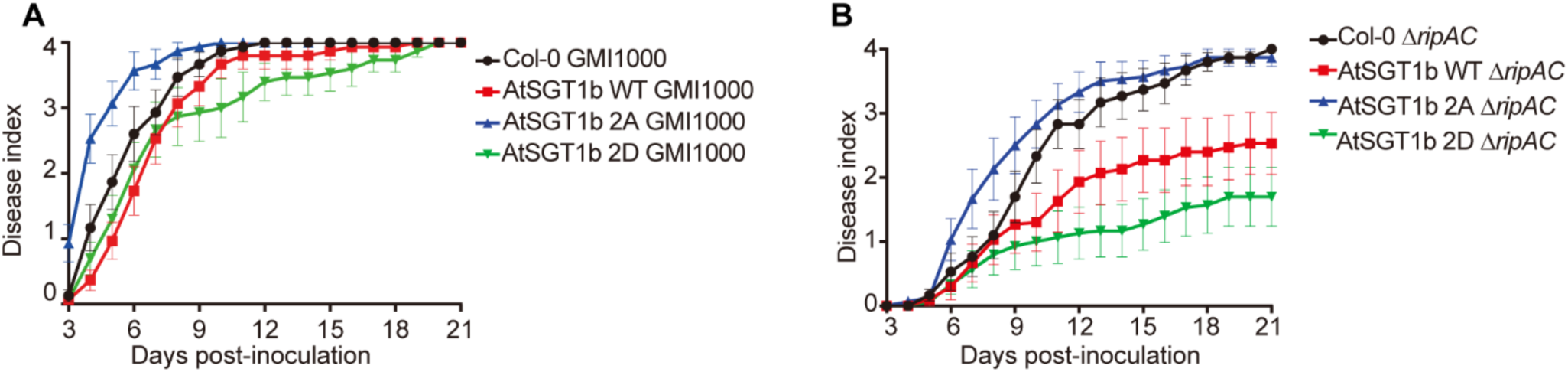
SGT1 phosphorylation contributes to resistance against *R. solanacearum* in Arabidopsis. Soil-drenching inoculation assays in Arabidopsis transgenic lines overexpressing AtSGT1b variants were performed with GMI1000 WT (A), or the Δ*ripAC* mutant (B). n=15 plants per genotype. The results are represented as disease progression, showing the average wilting symptoms in a scale from 0 to 4 (mean ± SEM). These experiments were repeated 3 times with similar results. See figure S13 for a composite representation.

## Discussion

Plant invasion by bacterial pathogens leads to the recognition of bacterial elicitors, either PAMPs or effectors, which, in turn, promotes the production and activation of additional immune receptors. Such feedback loop contributes to the robustness of the plant immune system. To achieve a successful infection and establish a compatible interaction, bacteria should evolve to suppress the activation of the immune responses triggered by the recognition of their PAMPs or effectors. Despite the importance of T3Es for the bacterial infection process, single effector knockout mutants often lack virulence phenotypes due to functional redundancy among effectors. This may be particularly important in strains belonging to the *R. solanacearum* species complex, where single strains may secrete more than 70 T3Es (Peeters et al., 2013). Although such large number of effectors contribute to the infection process, this also poses a risk for *R. solanacearum*, since each effector could potentially be recognized by intracellular NLRs, resulting in ETI. To maintain pathogenicity, bacterial pathogens need to adapt to new recognition events by losing these recognized effectors or evolving additional effectors that suppress ETI. This pathoadaptation becomes evident in the case of RipE1: a *R. solanacearum* GMI1000 derivative strain carrying mutations in *popP1* and *avrA* is pathogenic in *N. benthamiana* (Poueymiro et al., 2009), despite secreting RipE1, which activates ETI in this plant species (Sang et al., 2019). This suggests that *R. solanacearum* has evolved T3Es that suppress immunity triggered by RipE1 (and potentially other T3Es).

In this work, we found that the *R. solanacearum* T3E RipAC interacts with SGT1 (Figure 2), a major regulator of the activity of NLRs in the activation of ETI (Shirasu, 2009). SGT1 has been shown to regulate NLR homeostasis, either by contributing to protein stability or by mediating their degradation (Azevedo et al., 2006; Holt et al., 2005). RipAC was able to interact with a truncated SGT1 containing only the CS+SGS domains, but not with the CS domain alone, suggesting that RipAC interacts with the SGS domain, or that it requires both CS and SGS domains for interaction. The SGS domain is essential for SGT1 immune functions and mediates SGT1 interaction with the LRRs of NLRs and the co-chaperone HSC70 (Noël et al., 2007), although its biochemical function remains unclear.

We found that RipAC prevents the interaction between MAPKs and SGT1 in plant cells, leading to a reduced phosphorylation of SGT1 in the SGS domain (Figure 6), suggesting that MAPKs also interact with this domain. MAPKs have been shown to phosphorylate SGT1 from tobacco and maize in *N. benthamiana* (Hoser et al., 2013; Redkar et al., 2015), contributing to the hypothesis that MAPK-mediated phosphorylation of SGT1 may be important for its function. This notion is also supported by our data, showing that mutations that mimic phosphorylation in the observed SGT1 phosphosites lead to an enhanced activity of the NLR RPS2 and other SGT1-dependent ETI responses (Figures 5 and S10). Interestingly, the enhanced RPS2 activity caused by the T346D phospho-mimic mutation correlated with an enhanced dissociation between SGT1 (T346D) and RPS2 (Figure 5), suggesting that MAPK-mediated phosphorylation of SGT1 may contribute to NLR activation through the modulation of SGT1 interactions with NLRs (Figure S12). Considering that NLR activation subsequently enhances MAPK activation, this regulatory mechanism would represent a positive feedback loop between the activation of NLRs and MAPKs, mediated by the phosphorylation of SGT1 (Figure S12). Intriguingly, although RipAC suppresses SGT1-dependent immune responses, it does not seem to affect the accumulation of SGT1 or NLRs (in this case, RPS2), which resembles the phenotype observed upon overexpression of the SGS-interacting protein HSC70 (Noël et al., 2007). Moreover, a mutant including a premature stop codon at P347 in AtSGT1b (disrupting the MAPK phosphorylation motif that includes T346) does not affect SGT1 accumulation, but abolishes the immune-associated functions of SGT1 (Botër et al., 2007), highlighting the importance of the MAPK phosphorylation motif for SGT1 function. Altogether, our data suggest that, by associating with the SGS domain, RipAC compromises SGT1 interaction with MAPKs and the subsequent phosphorylation of specific sites in the SGS domain, compromising the activation of immunity without altering the accumulation of associated proteins (Figure S12).

We have recently shown that RipE1-triggered immunity requires SGT1 (Sang et al., 2019). In this work, we show that SGT1 phosphorylation contributes to RipE1-triggered cell death (Figure S10), and, accordingly, RipAC inhibits RipE1-triggered cell death (Figure 3). Altogether, this suggests that RipAC may contribute to *R. solanacearum* infection process by inhibiting SGT1-dependent immunity triggered by RipE1 and potentially other T3Es. Before, a *ΔripAC* mutant strain was reported to show reduced fitness, compared to GMI1000, in eggplant leaves (Macho et al., 2010). Here we show that an equivalent mutant has impaired virulence in Arabidopsis and tomato, and Arabidopsis transgenic plants expressing RipAC become more susceptible to *R. solanacearum* infection (Figure 1), indicating that RipAC plays a significant role in the *R. solanacearum* infection process. Furthermore, we show that the attenuation displayed by the *ΔripAC* mutant is partially rescued by the overexpression of AtSGT1b T346A (2A) (Figure 7), suggesting that SGT1 phosphorylation is an important target underlying RipAC virulence activity.

Although the contribution of SGT1 to the establishment of ETI is well known, genetic analysis of its contribution to disease resistance is challenging, due to the deleterious effects caused by the abolishment of SGT1 expression (Shirasu, 2009). Interestingly, overexpression of AtSGT1b did not have a strong impact on plant susceptibility against GMI1000 WT, but enhanced plant resistance against the *ΔripAC* mutant (Figure 7), indicating that the contribution of AtSGT1b overexpression is particularly evident in the absence of RipAC. It is noteworthy that overexpression of AtSGT1b T346D (2D) strongly enhanced plant resistance against the *ΔripAC* mutant (Figure 7), suggesting that, besides the effect of RipAC on SGT1 phosphorylation, there is an additive effect with other potential virulence activities of RipAC. Altogether, these data highlights the relevance of the SGT1 immunity node and SGT1 phosphorylation in the context of *R. solanacearum* infection.

The T3E AvrBsT from *Xanthomonas campestris* also interacts with SGT1 from pepper, and interferes with its phosphorylation by the PIK1 kinase in two serine residues (in the CS domain and variable region) different from the ones identified in our study (Kim et al., 2014). However, in that case, AvrBsT triggers SGT1-mediated immunity, and the identified SGT1 phosphorylation sites are required for the activation of AvrBsT-triggered HR (Kim et al., 2014). Similarly, a secreted effector from the fungal pathogen *Ustilago maydis* also interferes with the MAPK-mediated phosphorylation of SGT1 from maize in a monocot-specific phosphorylation site between TPR and CS domains, contributing to the development of disease-associated tumors (Redkar et al., 2015). This demonstrates that pathogens from different kingdoms and with different lifestyles have evolved to target different functions of SGT1 as a key regulator of disease resistance. However, some of these effectors are indeed perceived by the plant immune system, potentially as a consequence of the ‘guarding’ of SGT1 as an important regulator of plant immunity. On the contrary, RipAC seems to manipulate SGT1 for virulence purposes without being detected by the plant immune system, suggesting that these host plants have not yet evolved a perception system for this targeting. Together with other reports, our discovery of the molecular mechanisms underlying the RipAC-mediated suppression of SGT1 function may contribute to the engineering of functional SGT1 complexes that abolish bacterial manipulation or respond to it by activating immunity.

## Methods

The biological and chemical resources used in this study are summarized in the Table S1. Primers used in this study are summarized in the Table S2.

### Plant materials

Regarding *Arabidopsis thaliana*, Dexamethasone (Dex)-inducible *Arabidopsis thaliana GVG:AvrRpt2* (*Dex:AvrRpt2*) (McNellis et al., 1998) in ecotype Col-0 have been described previously. *GVG:AvrRpt2* (*Dex:AvrRpt2*) was used to cross with either RipAC line #3 or Col-0, and the resulting F1 progenies were used for experiments. In the experiments with Arabidopsis seedlings in 1/2 MS media, the seedlings were kept on 1/2 MS plates in a growth chamber (22°C, 16 h light/8 h dark, 100-150 mE m^−2^ s^−1^) for germination and growth for 5 days, then transferred to 1/2 MS liquid culture for additional 7 days. For *Pseudomonas syringae* and *Ralstonia solanacearum* infection assays, Arabidopsis plants were cultivated in jiffy pots (Jiffy International, Kristiansand, Norway) in a short day chamber (22°C, 10 h light/14 h dark photoperiod, 100-150 mE m^−2^ s^−1^, 65% humidity) for 4-5 weeks. After soil drenching inoculation, the plants were kept in a growth chamber under the following conditions: 75% humidity, 12 h light, 130 mE m^−2^ s^−1^, 27°C, and 12 h darkness, 26°C for disease symptom scoring.

*Nicotiana benthamiana* plants were grown at 22°C in a walk-in chamber under 16 h light/8 h dark cycle and a light intensity of 100-150 mE m^−2^ s^−1^.

Tomato plants (*Solanum lycopersicum* cv. Moneymaker) were cultivated in jiffy pots (Jiffy International, Kristiansand, Norway) in growth chambers under controlled conditions (25°C, 16 h light/8 h dark photoperiod, 130 mE m^−2^ s^−1^, 65% humidity) for 4 weeks. After soil drenching inoculation, the plants were kept in a growth chamber under the following conditions: 75% humidity, 12 h light, 130 mE m^−2^ s^−1^, 27°C, and 12 h darkness, 26°C for disease symptom scoring.

### Bacterial strains

*Pseudomonas syringae* pv. *tomato* (*Pto*) strains, including *Pto* containing an empty vector (EV), or a vector expressing AvrRpm1, AvrRpt2, or AvrRps4, or the type-three secretion system (T3SS)-defective mutant *Pto hrcC*^−^, were cultured overnight at 28 °C in LB medium containing 25 μg mL^−1^ rifampicin and 25 μg mL^−1^ kanamycin.

*Ralstonia solanacearum* strains, including the phylotype I reference strain GMI1000, GMI1000 Δ*ripAC^−^* mutant, and GMI1000 *ripAC^+^* complementation strains, were cultured overnight at 28°C in complete BG liquid medium (Plener et al., 2012).

*Agrobacterium tumefaciens* strain GV3101 with different constructs was grown at 28 °C on LB agar media with appropriate antibiotics. The concentration for each antibiotic is: 25 μg mL^−1^ rifampicin, 50 μg mL^−1^ gentamicin, 50 μg mL^−1^ kanamycin, 50 μg mL^−1^ spectinomycin.

### Generation of plasmid constructs, transgenic plants, and *R. solanacearum* mutant strains

The primers used to generate constructs in this work are presented in Table S2. To generate the RipAC-GFP construct, the RipAC coding region (RSp0875) in pDONR207 (a gift from Anne-Claire Cazale and Nemo Peeters, LIPM, Toulouse, France) was sub-cloned into pGWB505 via LR reaction, resulting in pGWB505-RipAC, in which the expression of the *RipAC-GFP* fusion is driven by a *CaMV 35S* promoter. The pGWB502-AtSGT1b variants (WT, 2A, 2D) and pGWB505-RipAC recombinant constructs were transformed into *A. tumefaciens* GV3101 and was then transformed in Arabidopsis Col-0 wild-type (WT) through the floral dipping method (Clough and Bent, 1998). Transgenic Arabidopsis plants were selected using hygromycin (50 μg mL^−1^) and were further confirmed by western blot using an anti-RipAC custom antibody. All the experiments using RipAC transgenic Arabidopsis were performed using two independent T4 homozygous lines.

To generate the *R. solanacearum* Δ*ripAC^−^* mutant, the *RipAC* coding region was replaced by gentamicin resistant gene using homologous recombination method (Zumaquero et al., 2010). The *RipAC* flanking regions, left border (LB) and right border (RB), were amplified by PCR and were recombined in the pEASYBLUNT vector, while the gentamicin resistance gene was inserted between the LB and RB by *Eco*R I digestion and T4 ligation. The resulting plasmid pEASYBLUNT-LB-Gm-RB was introduced into *R. solanacearum* GMI1000 WT strain by natural transformation (González et al., 2011). The Δ*ripAC^−^* mutant was selected using gentamicin (10 μg mL^−1^) and PCR using RipAC coding region specific primers (Table S2). To complement *RipAC* in the Δ*ripAC^−^* mutant, the *RipAC* promoter (423 bp upstream of ATG of the *RipABC* operon (Guéneron et al., 2000)) was cloned into pRCT-GWY (Henry et al., 2017), after which the *RipAC* coding region was introduced by LR reaction to result in a pRCT-pRipAC-RipAC construct. The integrative pRCT-pRipAC-RipAC plasmid, triggering the expression of RipAC under the control of the native *RipABC* operon promoter, was transformed into *R. solanacearum* Δ*ripAC^−^* mutant strain by natural transformation. The complementation strain *ripAC*^+^ was selected using Tetracycline (10 μg mL^−1^) and PCR using the RipAC coding region specific primers (Table S2).

### Pathogen inoculation assays

For *R. solanacearum* in soil inoculation, 15 four-to-five-week old Arabidopsis plants per genotype or 12 tomato plants for each bacterial strain (grown in Jiffy pots) were inoculated by soil drenching with a bacterial suspension containing 10^8^ colony-forming units per mL (CFU mL^−1^). 300 mL of inoculum of each strain was used to soak each treatment. After 20-minute incubation with the bacterial inoculum, plants were transferred from the bacterial solution to a bed of potting mixture soil in a new tray (Vailleau et al., 2007). Scoring of visual disease symptoms on the basis of a scale ranging from ‘0’ (no symptoms) to ‘4’ (complete wilting) was performed as previously described (Vailleau et al., 2007). To perform the survival analysis, the disease scoring was transformed into binary data with the criteria: the disease index lower than 2 was defined as ‘0’, while the disease index equal or higher than 2 was defined as ‘1’ in terms of the corresponding time (days post-inoculation, dpi) (Remigi et al., 2011). For stem injection assays, 5 μL of a 10^6^ CFU mL^−1^ bacterial suspension was injected into the stems of 4-week-old tomato plants and 2.5 μL xylem sap was collected from each plant for bacterial number quantification 3 dpi. Injections were performed 2 cm below the cotyledon emerging site in the stem, while the samples were taken at the cotyledon emerging site.

For *Pto* inoculation, different *Pto* strains were resuspended in water at 1×10^5^ CFU mL^−1^. The bacterial suspensions were then infiltrated into 4-5-week-old Arabidopsis leaves. Bacterial numbers were determined 3 dpi.

### Transient gene expression and ion leakage measurements

For protein accumulation, microsome fractionation, confocal microscopic observation, split-luciferase complementation (Split-LUC), and co-immunoprecipitation assays, *A. tumefaciens* GV3101 carrying different constructs were infiltrated into leaves of 5-week-old *N. benthamiana.* The OD600 used was 0.5 for each strain in all the assays, except for Split-LUC assays, for which we used OD600=0.2. To prepare the inoculum, *A. tumefaciens* was incubated in the infiltration buffer (10 mM MgCl_2_, 10 mM MES pH 5.6, and 150 μM acetosyringone) for 2 h. For effector-triggered immunity suppression assay, *A. tumefaciens* expressing RipAC-GFP or GFP were infiltrated into *N. benthamiana* leaves with an OD600=0.5, one day after which *A. tumefaciens* expressing the cell death-inducing agents were infiltrated at the same spots with an OD600=0.15. Pictures were taken 5-d-post expression of the cell death-inducing agent. Cell death was observed using a BIO-RAD GelDoc XR^+^ with Image Lab software.

For ion leakage measurement assays, the AtSGT1b variants were infiltrated at an OD600 of 0.5 and the cell death-inducing agents: 35S-RPS2-HA, Avr3a/R3a pair, RipE1, or BAX, was infiltrated at an OD600 of 0.15 in the same infiltrated spots one-day-post infiltration with *A. tumefaciens* expressing the AtSGT1b variants. Three leaf discs (diameter=10mm) were collected per infiltration site into 4 mL ddH_2_O in 12-well plate at 21-h-post infiltration with *A. tumefaciens* expressing cell death-inducing agents, then were placed on the bench top for 1 hour to remove the ion leakage caused by the wounding. Leaf discs were carefully transferred to fresh plates with ddH_2_O. Ion leakage was determined with ORION STAR A212 conductivity meter (Thermo Scientific) at the indicated time points. In the graph, 24 hpi indicates 24-hours post infiltration with *A. tumefaciens* expressing cell death-inducing agents.

### Protein extraction, microsome fractionation, and immunoblot analysis

To extract protein samples, 12-day-old Arabidopsis seedlings and leaf discs (diameter=18mm) from *N. benthamiana* were frozen in liquid nitrogen and ground with a Tissue Lyser (QIAGEN, Hilden, Nordrhein-Westfalen, Germany). Samples were subsequently homogenized in protein extraction buffer (100 mM Tris (pH 8), 150 mM NaCl, 10% Glycerol, 1% IGEPAL, 5 mM EDTA, 5 mM DTT, 1% Protease inhibitor cocktail, 2 mM PMSF, 10 mM sodium molybdate, 10 mM sodium fluoride, 2 mM sodium orthovanadate). To isolate the microsome fraction, proteins were extracted in buffer H (250 mM sucrose, 50 mM N-(2-hydroxyethyl) piperazine-N’-(2-ethanesulfonic acid) (HEPES)-KOH (pH 7.5), 5 % (v/v) glycerol, 50 mM sodium pyrophosphate decahydrate, 1 mM sodium molybdate dihydrate, 25 mM sodium fluoride, 10 mM EDTA, 0.5 % (w/v) polyvinylpyrrolidone (PVP-10) with a final concentration of 3 mM DTT and 1% protease inhibitor cocktail) and subjected to microsomal protein enrichment using differential centrifugation and Brij-58 treatment as previously described (Collins et al., 2017). The resulting protein samples were boiled at 70 °C for 10 minutes in Laemmli buffer and loaded in SDS-PAGE acrylamide gels for western blot. All the immunoblots were analyzed using appropriate antibodies as indicated in the figures. Molecular weight (kDa) marker bands are indicated for reference.

### MAP kinase activation assays

To measure the MAPK activation triggered by cell death inducers, *N. benthamiana* leaf discs were collected 24-h-post infiltration with *A. tumefaciens* expressing the cell death inducers. For ETI-triggered MAPK activation assays in Arabidopsis, four-week-old Col-0 plants were infiltrated with *Pto* EV, *Pto hrcC*^−^, *Pto* AvrRpt2, or *Pto* AvrRps4 (1×10^7^ CFU mL^−1^) and samples were collected at the indicated time points as previously described (Su et al., 2018; Tsuda et al., 2013). To evaluate the MAPK activation in *Dex:AvrRpt2* Arabidopsis, 12-d-old *Dex:AvrRpt2/*Col-0 or *Dex:AvrRpt2/*AC #3 F1 crossing seedlings grown in liquid media were treated with 30 μM Dex for different times as indicated and previously described (Guan et al., 2015; Kadota et al., 2019). After protein extraction, the protein samples were separated in 10% SDS-PAGE gels and the western blots were probed with anti pMAPK antibodies to determine MAPK activation as previously described (Macho et al., 2012). Detection of SGT1 accumulation and SGT1 phosphorylation were also performed with the same samples using custom anti-SGT1 and anti-pSGT1 antibodies. The induction of AvrRpt2 was determined by RT-PCR. Blots were stained with Coomassie Brilliant Blue (CBB) or were probed with anti-actin to verify equal loading.

### Large-scale immunoprecipitation and LC-MS/MS analysis

Large-scale immunoprecipitation assays for LC-MS/MS analysis were performed as previously described with several modifications (Sang et al., 2018). Two grams of *N. benthamiana* leaf tissues were collected at 2 days after infiltration with *A. tumefaciens* and frozen in liquid nitrogen. The protein extracts were homogenized in protein extraction buffer (100 mM Tris-HCl pH8, 150 mM NaCl, 10% glycerol, 5 mM EDTA, 10 mM DTT, 2 mM PMSF, 10 mM NaF, 10 mM Na_2_MoO_4_, 2 mM NaVO_3_, 1%(v/v) NP-40, 1%(v/v) plant protease inhibitor cocktail (Sigma). The protein extracts were cleaned by 2 rounds of 10 min x 15, 000 g centrifugation to remove the tissue debris. 30 μL GFP-trap beads (ChromoTek, Germany) or ANTI-FLAG M2 Affinity Agarose Gel (Sigma) was added to the clean protein extract and incubated for one hour at 4 °C with an end-to-end slow but constant rotation. GFP-Trap beads or ANTI-FLAG beads were washed four times with 1 mL cold wash buffer (100 mM Tris-HCl pH 8, 150 mM NaCl, 10% glycerol, 2 mM DTT, 10 mM NaF, 10 mM Na_2_MoO_4_, 2 mM NaVO_3_, 1%(v/v) plant protease inhibitor cocktail (Sigma), 0.5%(v/v) NP-40 for GFP-Trap beads, no NP-40 for anti-FLAG M2 beads) and the washed beads were subjected to Mass Spectrometric analysis as previously described to identify interacting proteins (Sang et al., 2018) and phosphorylated peptides (Wei et al., 2017).

### Co-immunoprecipitation

One gram of *N. benthamiana* leaf tissues was collected at 2 days after infiltration with *A. tumefaciens* and frozen in liquid nitrogen. Total proteins were extracted as indicated above and immunoprecipitation was performed with 15 μL of GFP-trap beads (ChromoTek, Germany) or ANTI-FLAG M2 Affinity Agarose Gel (Sigma) as described above. Beads were washed 4 times with wash buffer with different detergent concentrations. To specify, in Figures 2A and S5F, there is no detergent in the wash buffer, while in Figures 3A and 4G, the concentration for detergent in wash buffer is 0.2%. The proteins were stripped from the beads by boiling in 40 µL Laemmli buffer for 10 minutes at 70 °C. The immunoprecipitated proteins were separated on SDS-PAGE gels for western blot analysis with the indicated antibodies. Blots were stained with CBB to verify equal loading.

### Split-LUC assays

Split-LUC assays were performed as previously described (Chen et al., 2008; Wang et al., 2018). Briefly, *A. tumefaciens* strains containing the desired plasmids were hand-infiltrated into *N. benthamiana* leaves. Split-LUC assays were performed both qualitatively and quantitatively after 2 dpi or 1dpi, respectively, in the RPS2 expression assays. For the CCD imaging, the leaves were infiltrated with 0.5 mM luciferin in water and kept in the dark for 5 min before CCD imaging. The images were taken with either Lumazone 1300B (Scientific Instrument, West Palm Beach, FL, US) or NightShade LB 985 (Berthold, Bad Wildbad, Germany). To quantify the luciferase signal, leaf discs (diameter=4 mm) were collected into a 96-well microplate (PerkinElmer, Waltham, MA, US) with 100 μL H_2_O. Then the leaf discs were incubated with 100 μL water containing 0.5 mM luciferin in a 96-well plate wrapped with foil paper to remove the background luminescence for 5 min, and the luminescence was recorded with a Microplate luminescence reader (Varioskan flash, Thermo Scientific, USA). Each data point contains at least eight replicates. The protein accumulation was determined by immunoblot as described above.

### Sequence analysis

The SGT1 protein sequences were retrieved from Phytozome (https://phytozome.jgi.doe.gov/pz/portal.html) and sequence alignments were generated using Clustal Omega (https://www.ebi.ac.uk/Tools/msa/clustalo/) with default settings. The consensus MAPK-mediated phosphorylation sites were generated using WebLogo (https://weblogo.berkeley.edu).

### Confocal microscopy

To determine the protein subcellular localization, bimolecular fluorescence complementation (BiFC), and protein co-localization in *N. benthamiana*, leaf discs were collected 2 dpi and observed using a Leica TCS SP8 (Leica, Mannheim, Germany) confocal microscope with the following excitation wavelengths: GFP, 488nm; YFP, 514nm; RFP, 561 nm (Rosas-Diaz et al., 2018).

### Protoplast assays

Protoplast transient expression assays were performed as described previously (Cui et al., 2013; Yoo et al., 2007). In brief, 200 μL of protoplasts (2 x 10^6^ cells) were mixed with plasmids (20 μg for each construct) with a 1:1 plasmid ratio for every combination. The protoplasts were incubated at room temperature overnight and were harvested by centrifugation at 100 g for 2 min. After removing the supernatant, 90 μL protein extraction buffer was added to the protoplasts. After boiling in the Laemmli buffer, the protein samples were subjected to western blot analysis.

### Generation of custom antibodies

To raise anti-RipAC antibodies, purified recombinant GST-RipAC_241-540aa_ expressed in *E. coli* was used as antigen. The polyclonal antiserum from immunized rabbits was purified by affinity chromatography using recombinant GST-RipAC_241-540aa_, and the eluate was used as anti-RipAC antibody (Abclonal Co., Wuhan, China). The anti-RipAC antibody specificity was determined by using RipAC-GFP transient expression in *N. benthamiana*. The SGT1 phosphosite-specific antibodies (anti-AtSGT1b pT346) were generated by Abclonal Co. Briefly, the synthesized phospho-peptide [ES(T-p)PPDGME-C] was conjugated to keyhole limpet hemocyanin (KLH) carrier to immunize rabbits. The rabbit polyclonal antiserum was purified by affinity chromatography using phospho-peptide and the eluate was cleaned by passing through the column coupled with control synthesized-peptide (ESTPPDGME-C) to remove the non-specific antibodies. To determine the pSGT1 antibody specificity, an *in vitro* ELISA and an *in vivo* Calf Intestinal Alkaline Phosphatase (CIAP) treatment were performed. In the ELISA assay, the phospho-peptide and control peptide were spotted on the nitrocellulose filter membrane and probed with different diluted anti-pSGT1 antibody. In the CIAP treatment, protein samples from two 12-d-old Col-0 seedlings were extracted with 300 μL 1 x NEB CutSmart buffer (50 mM Potassium Acetate, 20 mM Tris-acetate, 10 mM Magnesium Acetate, 100 µg/ml BSA, pH 7.9) with 1% protease inhibitor cocktail. 2 μL CIAP enzyme (New England Lab) was added to a vial of 40 μL protein extract, while another 40 μL protein extract with no CIAP was used as control. The treatment was performed by incubating protein extract at 37 °C for 1h, after which the protein sample was subjected to western blot analysis with different antibodies. The anti-pSGT1 antibody was used to determine the effect of CIAP treatment, while the anti-SGT1 antibody was used to show the endogenous SGT1 abundance.

### Quantification and statistical analysis

Statistical analyses were performed with Prism 7 software (GraphPad). The data are presented as mean ± SEM. The statistical analysis methods are described in the figure legends.

## Author contributions

G.Y. performed most of the experiments and analyzed data; L.X. initiated the project and performed experiments; H.X., W.Y., and J.S.R. performed experiments; Y.S., R.J.L.M, and Y.W. provided new techniques and feedback; A.P.M. conceived and supervised the project; G.Y and A.P.M. designed the experiments and wrote the manuscript with inputs from all the authors.

## Acknowledgements

We thank Nemo Peeters and Anne-Claire Cazale for sharing unpublished biological materials, Yasuhiro Kadota and Ken Shirasu for helpful discussions and sharing biological materials, Suomeng Dong, Jian-Min Zhou, Gitta Coaker, and Laurent Deslandes for sharing biological materials, Rosa Lozano-Duran for critical reading of this manuscript, and Xinyu Jian for technical and administrative assistance during this work. We thank the PSC Cell Biology and Proteomics core facilities for assistance with confocal microscopy and LC-MS/MS analysis, respectively. This work was supported by the Strategic Priority Research Program of the Chinese Academy of Sciences (grant XDB27040204), the National Natural Science Foundation of China (NSFC; grant 31571973), the Chinese 1000 Talents Program, and the Shanghai Center for Plant Stress Biology (Chinese Academy of Sciences). Gang Yu was partially supported by the China Postdoctoral Science Foundation (grant 2016M600339). The authors have no conflict of interest to declare.

---

**Supplemental Figure 1.**
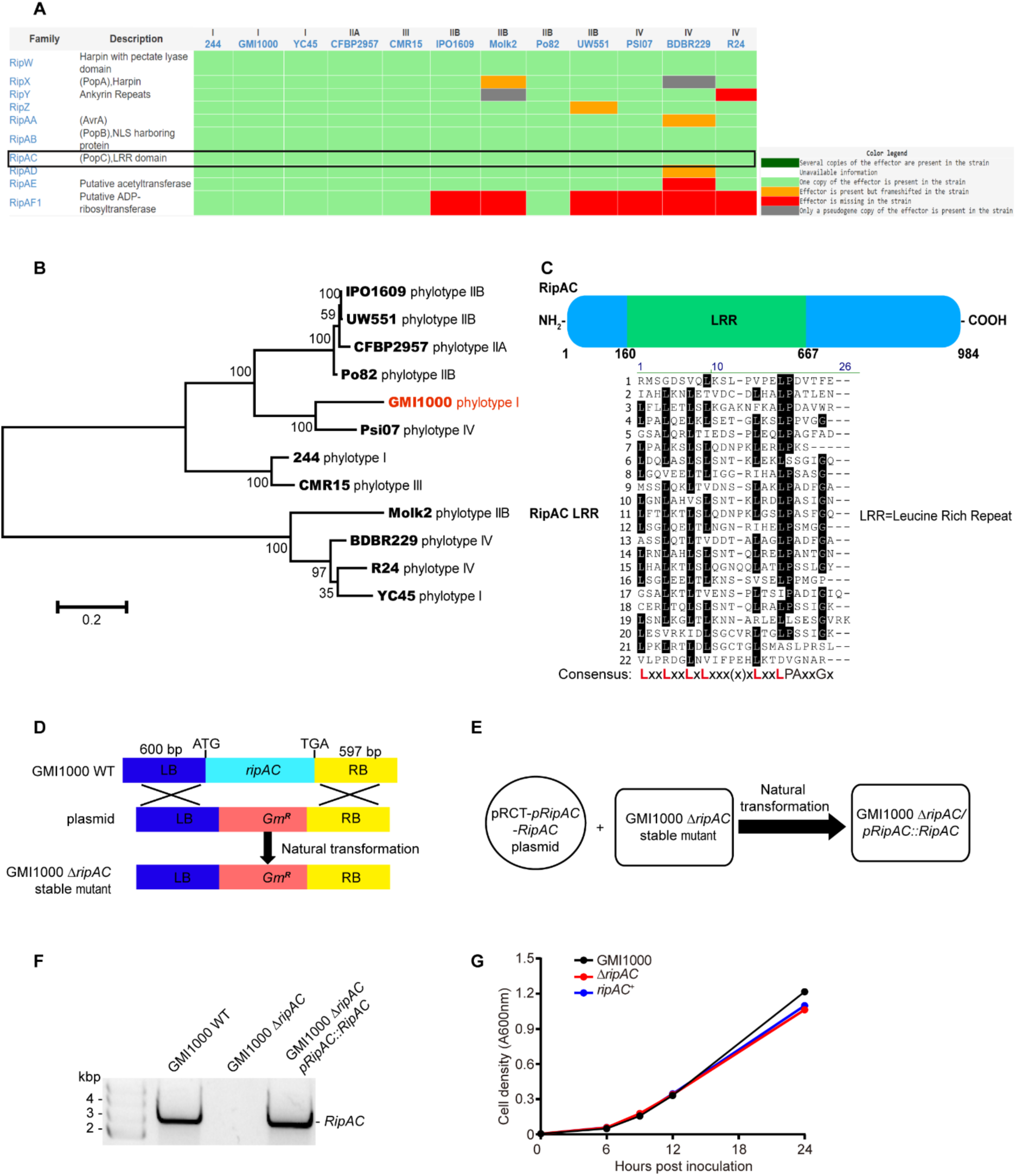
RipAC encodes a leucine rich repeat protein in *Ralstonia solanacearum*. (A) RipAC is a core effector present in most sequenced *R. solanacearum* strains. The data is retrieved from the Ralsto T3E website (https://iant.toulouse.inra.fr/bacteria/annotation/site/prj/T3Ev3/). (B) Phylogenetic analysis of RipAC proteins from different sequenced *R. solanacearum* strains. The phylogenetic tree was generated using the Maximum Likelihood method based on the JTT matrix-based model. The tree is drawn to scale with branch lengths meaning the measurement number of substitutions per site. (C) RipAC encodes a leucine rich repeat protein with no predicted enzymatic domain. The upper panel is the diagram of the RipAC protein with LRR domain; the lower panel is the detailed analysis of the RipAC LRR domain, and the numbers on the left indicate LRR numbers. (D) Diagram of the process of generation of the *R. solanacearum ΔripAC* mutant. The *ΔripAC* mutant was generated by homologous recombination method using a pEASYBLUNT-based plasmid as described in the methods section. (E) Generation of *RipAC* complementation strain in the *ΔripAC* mutant strain. A 423bp DNA fragment upstream of ATG of the *RipABC* operon was amplified and inserted into pRCT plasmid, and the RipAC coding region was shifted into the pRCT plasmid by LR reaction to result in the pRCT-*pRipAC*-*RipAC* expression cassette. The integrative pRCT-pRipAC-RipAC plasmid was mobilized into the *ΔripAC* mutant by natural transformation to result in *ripAC^+^*. (F) PCR characterization of the presence of the *RipAC* DNA fragment in different strains. A pair of PCR primers was designed to amplify the *RipAC* full-length coding region and the PCR was performed to examine the presence of *RipAC* gene. (G) Bacterial growth in nutrient-rich medium. GMI1000 WT, *ΔripAC*, and *ripAC^+^* strains were inoculated into the complete BG liquid medium with initial OD600=0.005 and the bacterial growth was monitored at the indicated time points measuring OD600 (mean ± SEM, n=3).

**Supplemental Figure 2.**
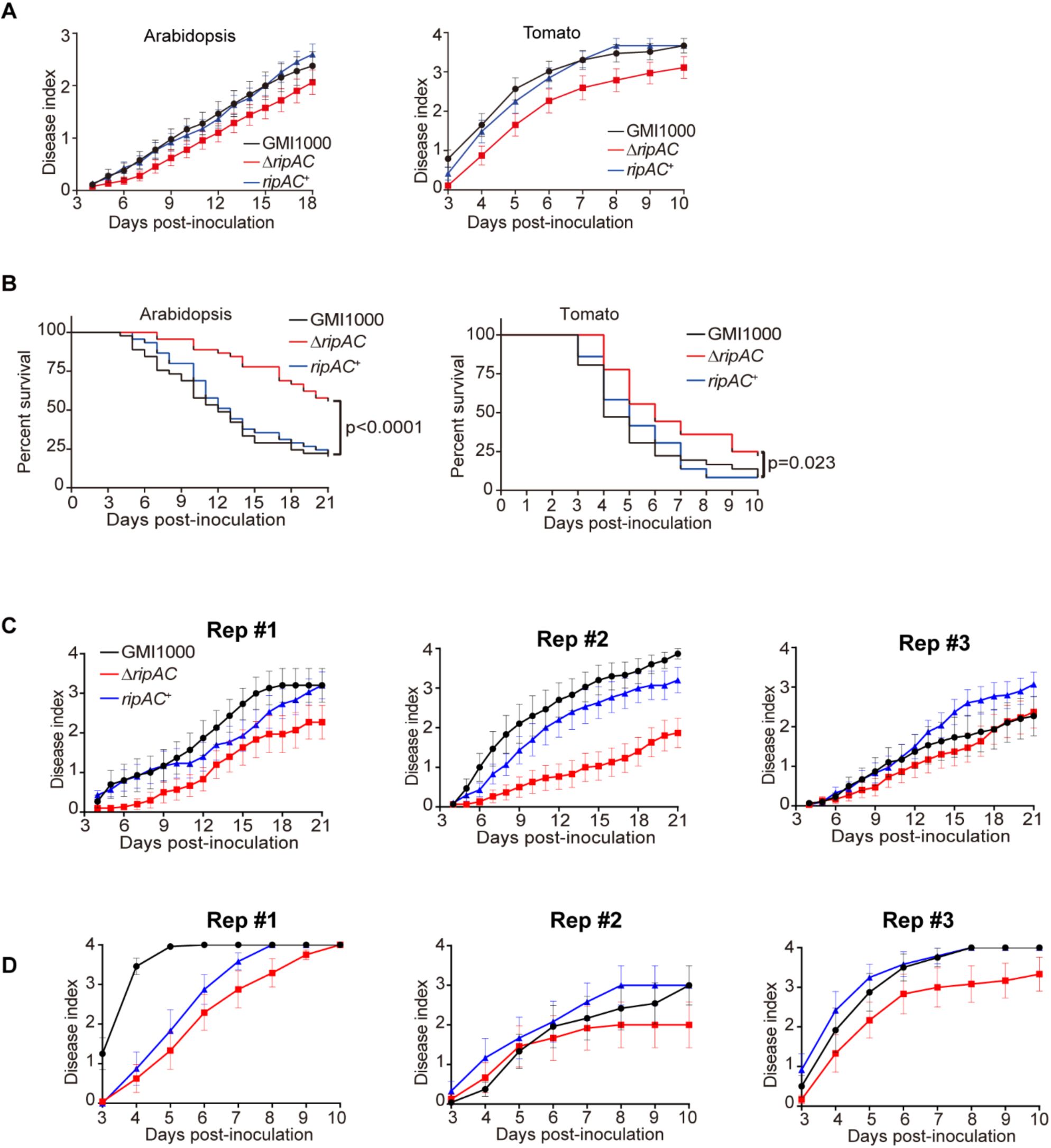
*Ralstonia solanacearum* inoculation assays in Arabidopsis and tomato. (A) Soil-drenching inoculation assays in Arabidopsis and tomato were performed with GMI1000 WT, Δ*ripAC* mutant, and RipAC complementation (*ripAC*^+^) strains. The results are represented as disease progression, showing the average wilting symptoms in a scale from 0 to 4. Each data point represents the mean disease index for three independent experiments (n = 45 for each strain in Arabidopsis assays or n = 36 for each strain in tomato assays). Each vertical bar represents the SEM from three independent experiments. (B) Survival analysis of Arabidopsis in (A). Statistical analysis were performed with Log-rank (Mantel-Cox) test (n = 45 for each strain in Arabidopsis assays or n = 36 for each strain in tomato assays). The calculated P values are presented in the graphs. (C) Three independent biological replicates of *R. solanacearum* soil drenching inoculation assays on Arabidopsis (Col-0 WT). (D) Three independent biological replicates of *R. solanacearum* soil drenching inoculation assays on tomato (cv. moneymaker). The soil drenching inoculation assays were performed with GMI1000 WT, *ΔripAC* mutant, and RipAC complementation (*ripAC*^+^) strains. In (C) and (D) N=15 plants per genotype (for Arabidopsis) or n=12 plants per genotype (for Tomato). The results are represented as disease progression, showing the average wilting symptoms in a scale from 0 to 4 (mean ± SEM).

**Supplemental Figure 3.**
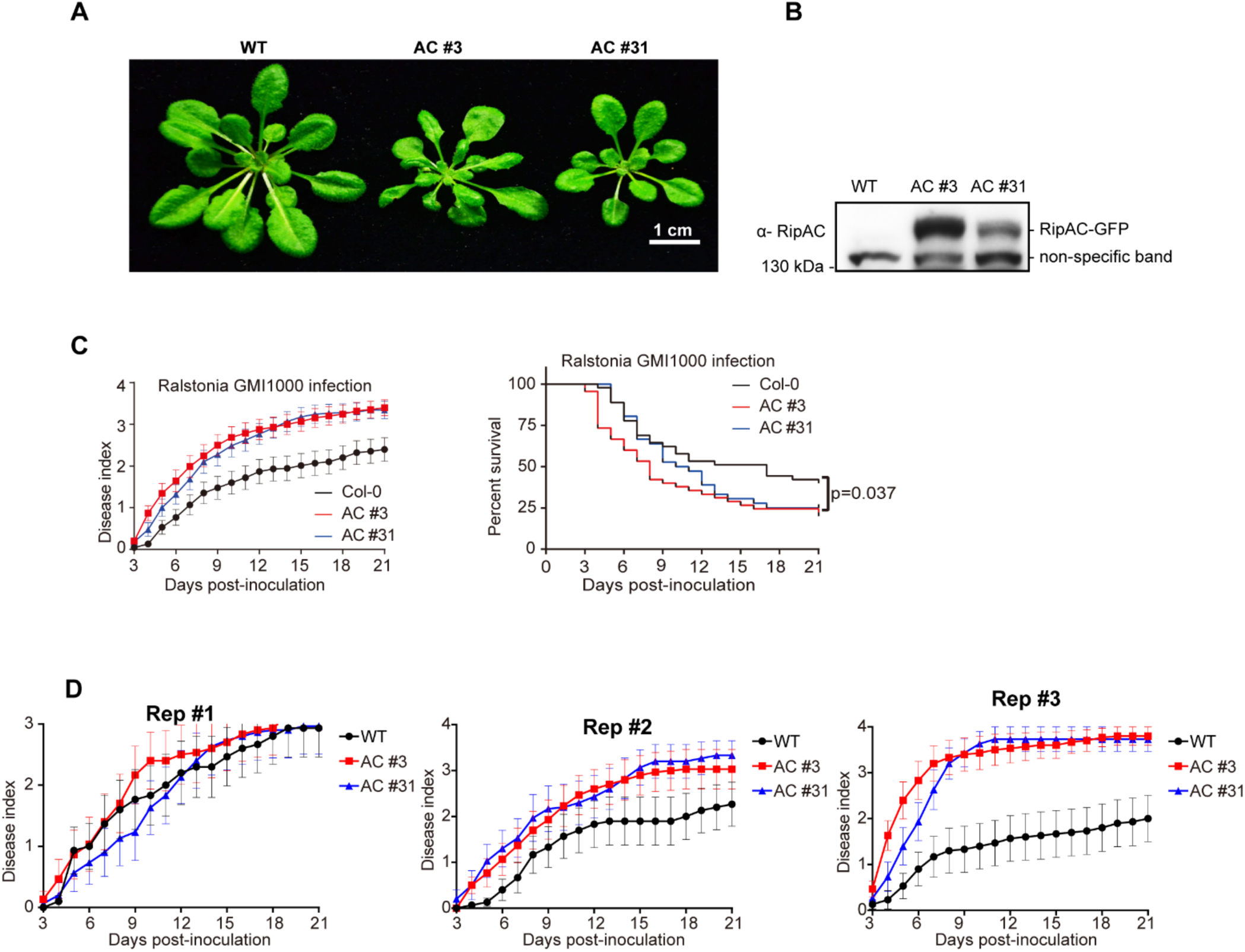
RipAC-GFP transgenic Arabidopsis are more susceptible to *R. solanacearum* GMI1000. (A) Typical developmental phenotypes of RipAC-GFP transgenic Arabidopsis. AC #3 and AC #31 are two independent transgenic lines (T4 generation). The picture shows 1-month-old Arabidopsis grown in a short-day growth chamber. (B) Western blot shows RipAC-GFP protein accumulation in transgenic Arabidopsis. Samples were taken at 12 days after germination. Blots were probed with antibody Anti-RipAC (1:5000). (C) Soil-drenching inoculation assays in RipAC-GFP transgenic lines with GMI1000 WT strain. The results are represented as disease progression, showing the average wilting symptoms in a scale from 0 to 4. Each data point represents the mean disease index for three independent experiments (n = 45 in total for different Arabidopsis genotypes). Each vertical bar represents the SEM from three independent experiments. Statistical analysis on the right graph was performed with Log-rank (Mantel-Cox) test (n = 45 in total for different Arabidopsis genotypes). The calculated P value is presented in the graph. (D) Three independent biological replicates of *R. solanacearum* soil drenching inoculation assays on Col-0 WT, AC #3 and AC #31. The in soil drenching inoculation assays were performed with the GMI1000 WT strain. n=15 plants per genotype. The results are represented as disease progression, showing the average wilting symptoms in a scale from 0 to 4 (mean ± SEM).

**Supplemental Figure 4.**
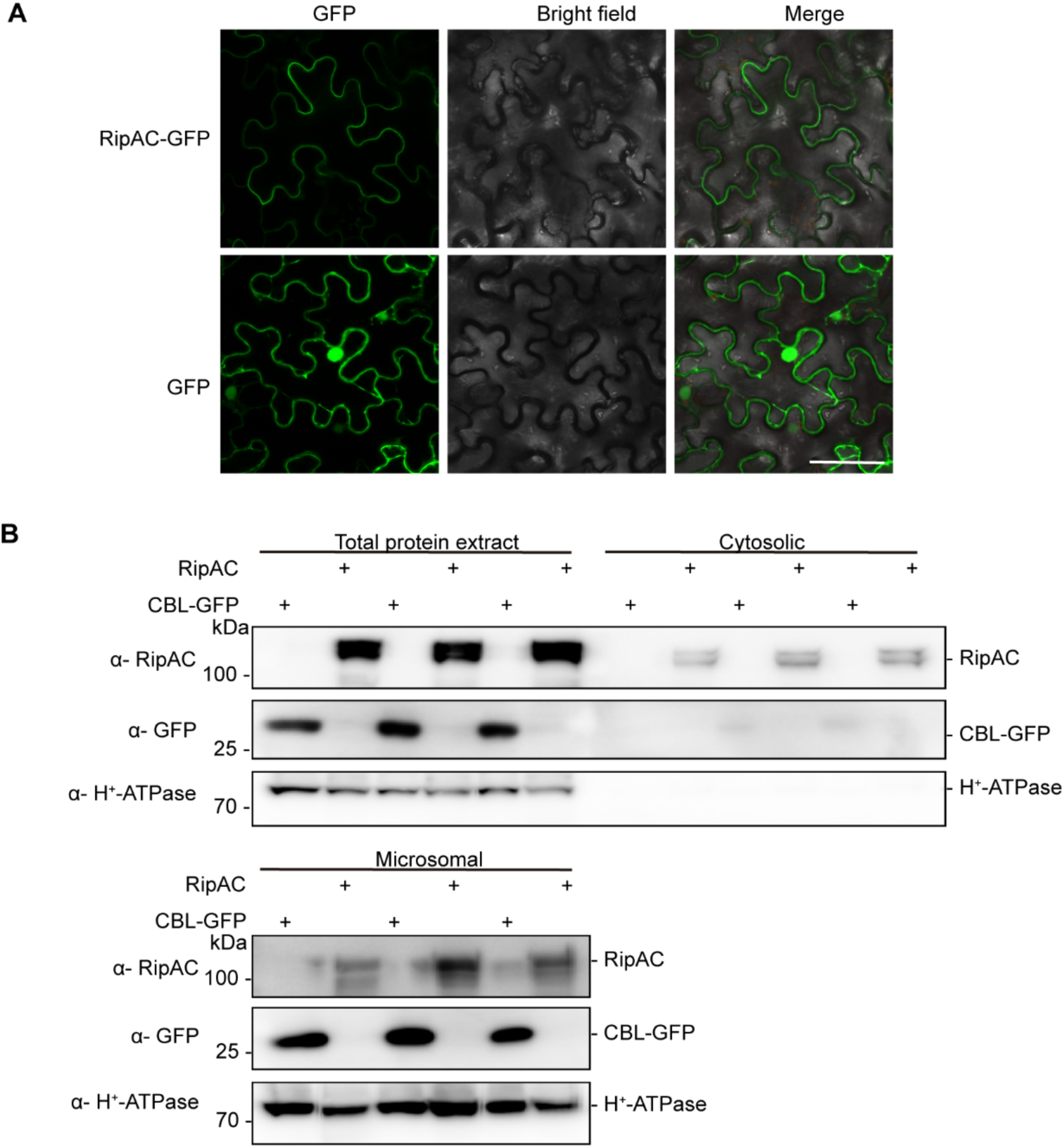
Subcellular localization of RipAC-GFP. (A) Subcellular localization of RipAC-GFP in plant cells. RipAC-GFP or free GFP were transiently expressed in *N. benthamiana* leaves using *Agrobacterium tumefaciens*, and the GFP fluorescence signal was observed 48 hpi using confocal microscopy. Scale bar=50 µm. (B) Microsome fractionation in *N. benthamiana*. Agrobacterium carrying RipAC or CBL-GFP was infiltrated into 5-week-old *N. benthamiana* leaves and samples were taken at 2dpi and then subjected to microsome fractionation. The total protein extraction was separated into the cytosolic fraction and the microsomal fraction using centrifugation as described in the methods section. Protein samples from total extract, cytosolic, and microsome fraction were used for western blot. The plasma-membrane protein H^+^-ATPase was used as a microsomal protein marker. Western blots from 3 biological replicates are represented.

**Supplemental Figure 5.**
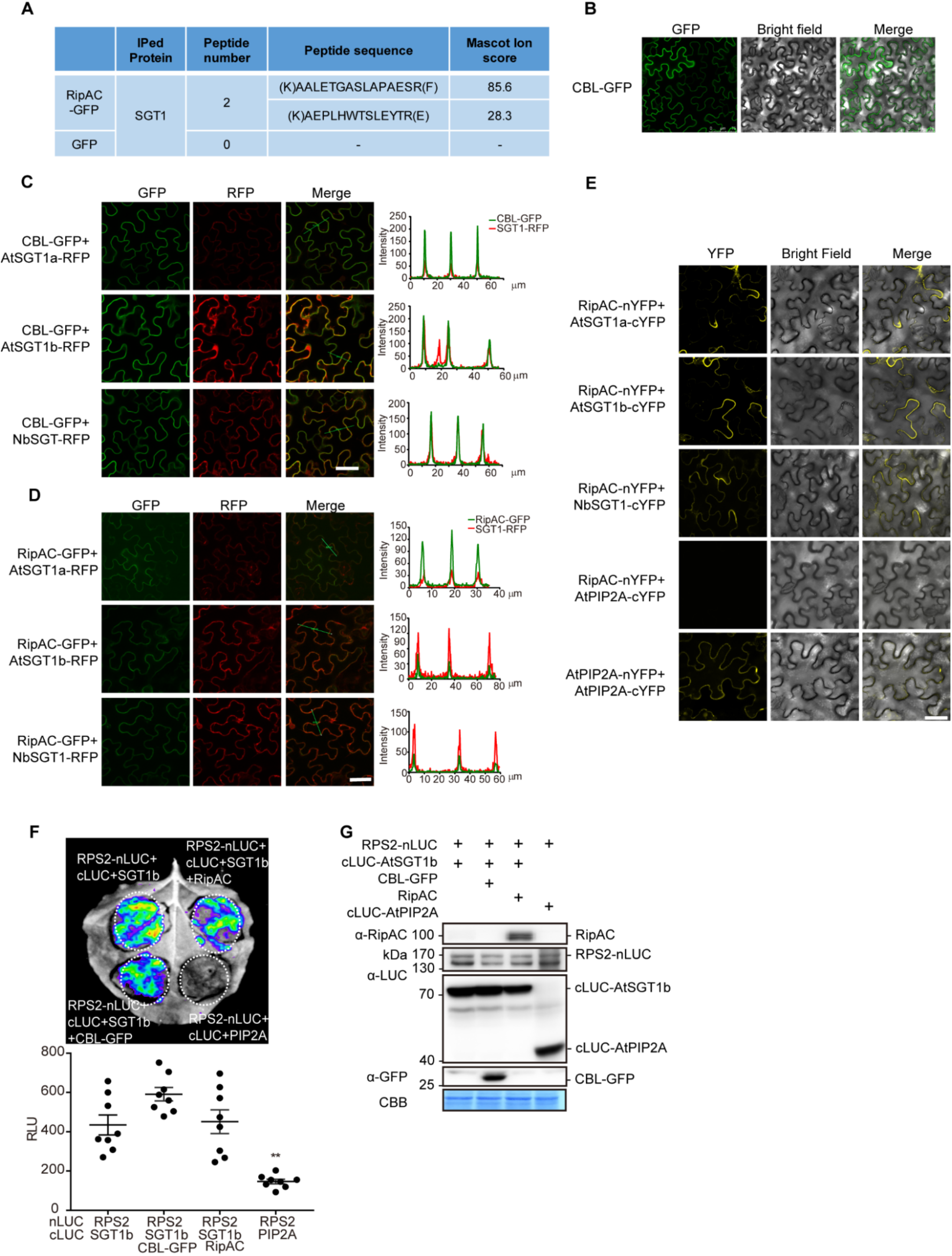
RipAC associates with SGT1s in plant cells. (A) RipAC-GFP or free GFP were transiently expressed in *N. benthamiana* leaves. The figure shows the unique NbSGT1 peptides identified exclusively in RipAC-GFP sample upon GFP immunoprecipitation followed by IP-MS/MS analysis. (B) CBL-GFP localizes at plasma membrane. bar=75 µm. (C) Co-localization analyses of plasma membrane-associated CBL-GFP protein and SGT1-RFP. (D) RipAC-GFP co-localizes with SGT1-RFP in plant cells. In (B), (C) or (D) CBL-GFP alone, CBL-GFP with SGT1-RFP (AtSGT1a, AtSGT1b, and NbSGT1) or RipAC-GFP with SGT1-RFP (AtSGT1a, AtSGT1b, and NbSGT1) was transiently expressed using Agrobacterium in *N. benthamiana*, and the GFP fluorescence signal was observed 48 hpi using confocal microscopy. In (C) and (D), the images show the fluorescence from GFP, RFP channel, and the merged fluorescence from both channels. The corresponding fluorescence intensity profiles (GFP, green; RFP, red) across the green lines are shown. Scale bar=50 µm. (E) Split-YFP complementation assay to determine direct interaction between RipAC and SGT1 in plant cells. The self-association of aquaporin AtPIP2A was used as a positive interaction control, while the RipAC-AtPIP2A combination was used as a negative control. Fluorescence signal was observed 48 hpi using confocal microscopy. Scale bar=50 µm. (F) Competitive Split-LUC showing that RipAC does not interfere with RPS2-SGT1b association. (G) Western blot shows the protein accumulation in (F). The RPS2-PIP2A combination was used as negative control. In all the competitive interaction assays, in addition to the interaction pair, RipAC or CBL-GFP (as negative control) were expressed to determine interference. In (F) luciferase activity was determined both qualitatively (CCD camera, higher panel) and quantitatively (microplate luminescence reader, lower panel). All the experiments were performed 3 times with similar results.

**Supplemental Figure 6.**
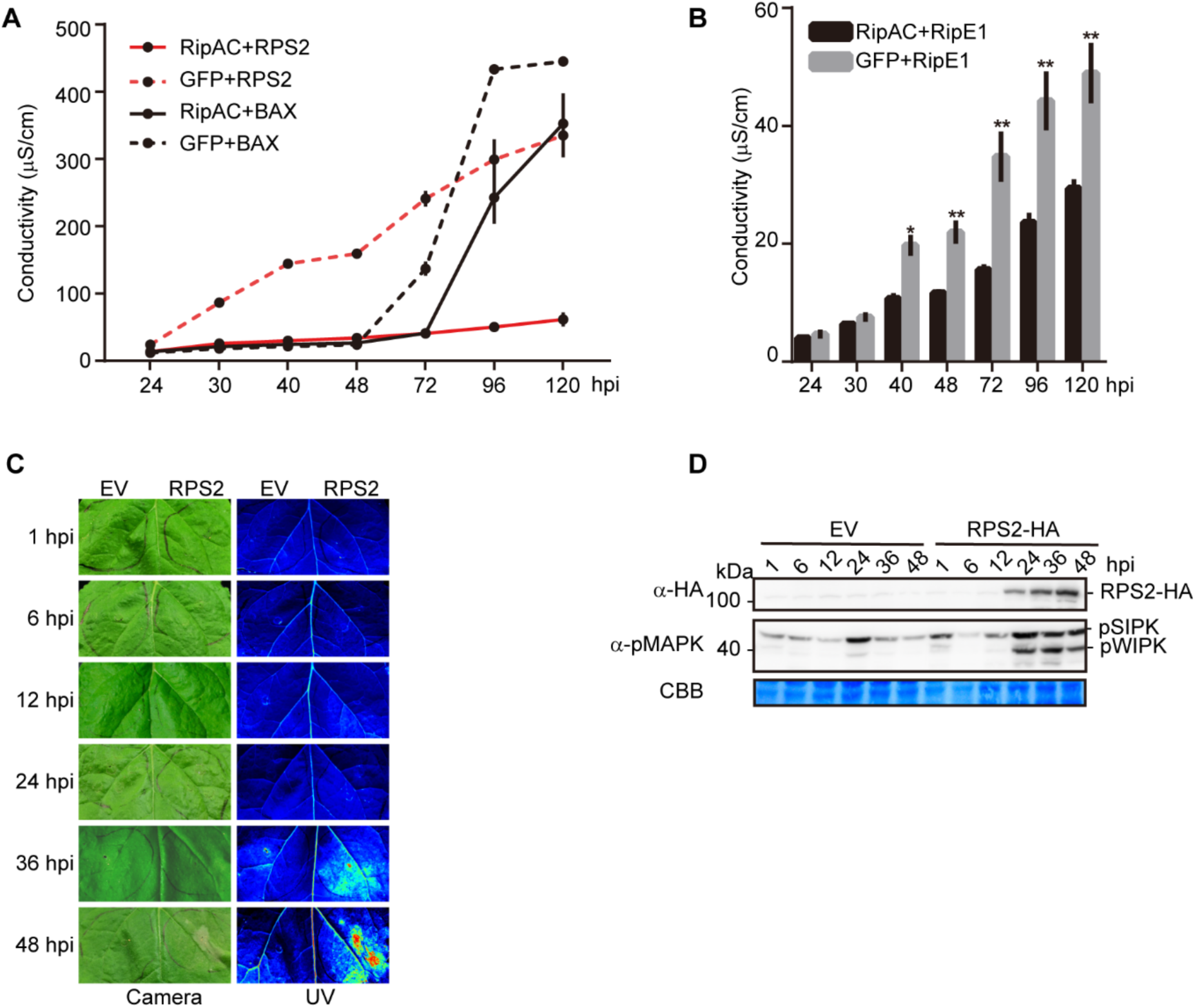
RipAC inhibits SGT1-dependent immune responses. (A) Ion leakage assays showing RipAC specifically suppresses RPS2-, but not BAX-, mediated cell death in *Nicotiana benthamiana*. Agrobacterium expressing RipAC or the GFP control (OD600=0.5) were infiltrated into *N. benthamiana* leaves 1 day before infiltration with Agrobacterium expressing RPS2 or BAX (OD600=0.15). Leaf discs were taken 21 hpi for conductivity measurements at the indicated time points. The time points in the x-axis are indicated as hpi with Agrobacterium expressing RPS2 or BAX (mean ± SEM, n=3, 4 replicates). (B) Ion leakage assays showing RipAC suppresses RipE1-mediated cell death in *N. benthamiana*. The ion leakage assays were performed the same as in (A) (mean ± SEM, n=3, * p<0.05, ** p<0.01, *t*-test, 3 replicates). (C) RPS2-triggered cell death in *Nicotiana benthamiana* was monitored by visible light and UV light. (D) Phosphorylation of NbSIPK and NbWIPK was detected using anti-pMAPK antibody.

**Supplemental Figure 7.**
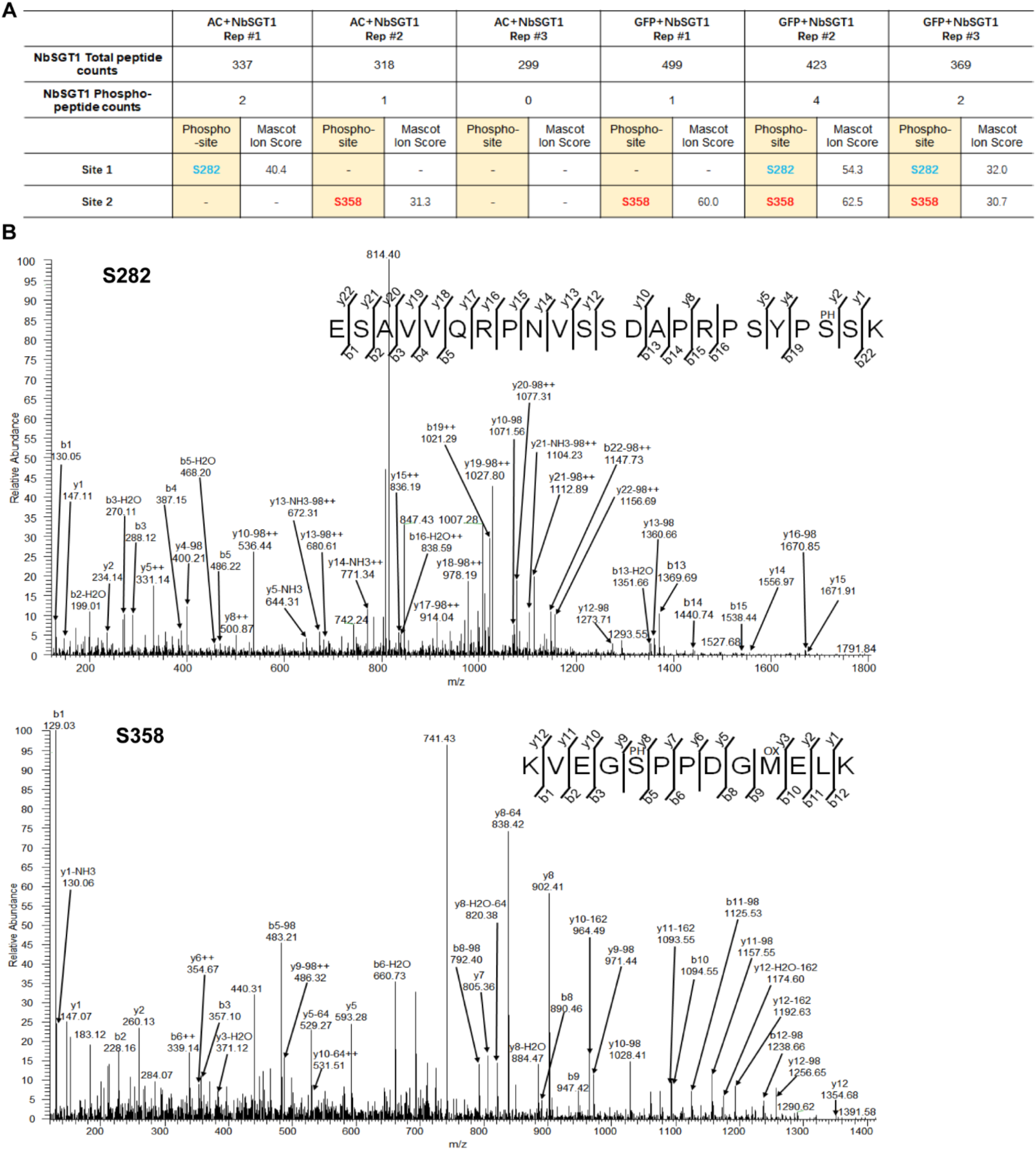
NbSGT1 phosphorylation in plant cells. (A) NbSGT1 is phosphorylated in *Nicotiana benthamiana*. Agrobacterium containing NbSGT1-FLAG with RipAC or GFP was infiltrated into 4-5 weeks old *N. benthamiana* plants and samples were harvested 48 hpi and were subjected to anti-FLAG IP-MS/MS. The phosphorylation of S282 and S358 is summarized from three biological IP-MS/MS replicates. (B) Representative MS/MS spectra showing phosphorylation of Ser282 and Ser358 in NbSGT1 expressed in *N. benthamiana*.

**Supplemental Figure 8.**
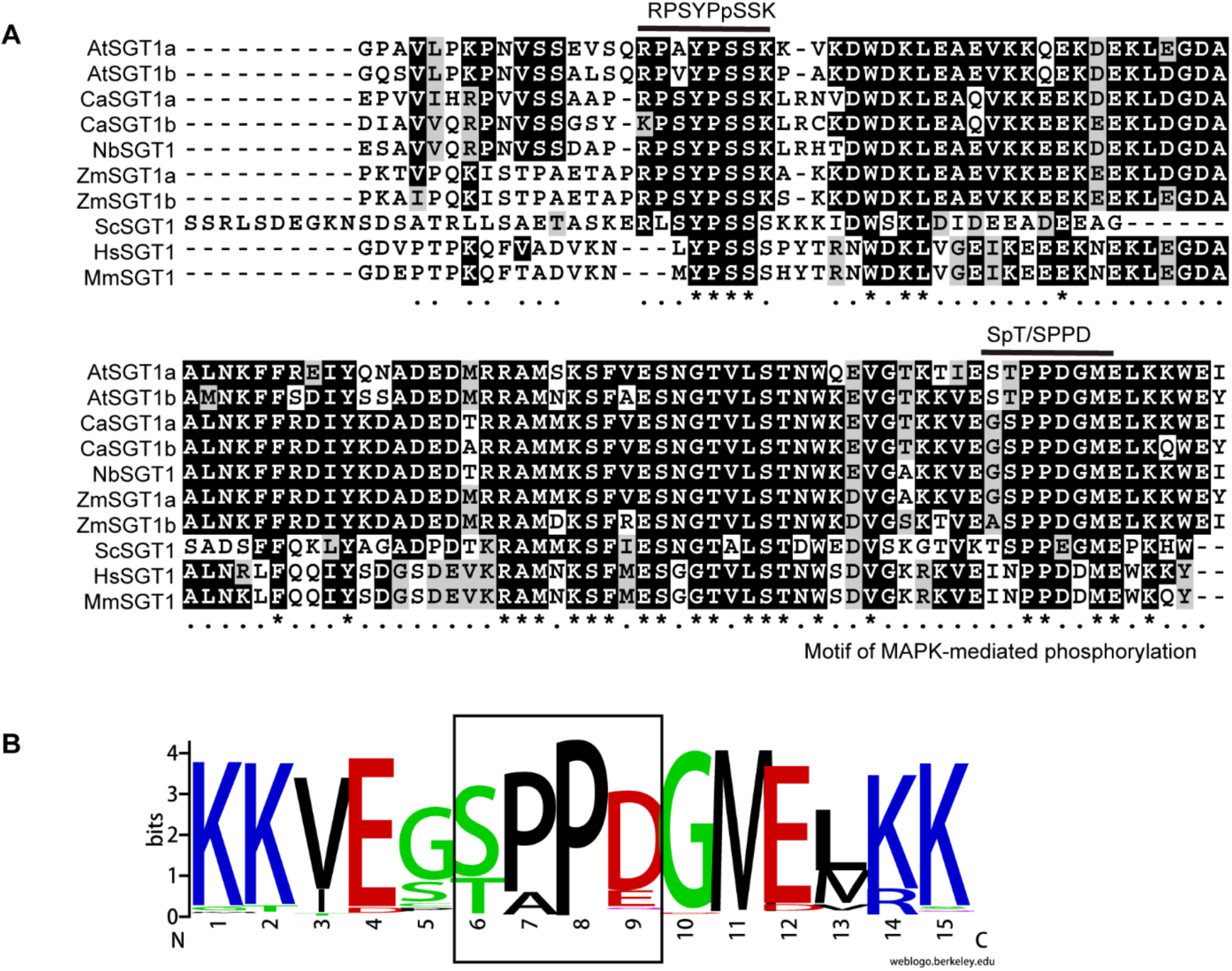
MAPK-mediated phosphorylation of SGT1 is conserved in plant kingdom. (A) Phosphorylation of S282 and S358 in NbSGT1 are located in the SGS (SGT1-specific) domain. The S358 is predicted to be a MAPK phosphorylation site. At = *Arabidopsis thaliana*; Ca= *Capsicum annuum*; Nb = *Nicotiana benthamiana*; Zm =*Zea mays*; Sc = *Saccharomyces cerevisiae*; Hs = *Homo sapiens*; Ms = *Mus musculus*. (B) MAPK-mediated phosphorylation motif in SGT1 proteins is conserved in the plant kingdom. SGT1 protein sequences from different plant species genome were retrieved from Phytozome (https://phytozome.jgi.doe.gov/pz/portal.html). Then the sequences were aligned with MEGA program and the MAPK-mediated phosphorylation motif “S/TP” were selected for Weblogo.

**Supplemental Figure 9.**
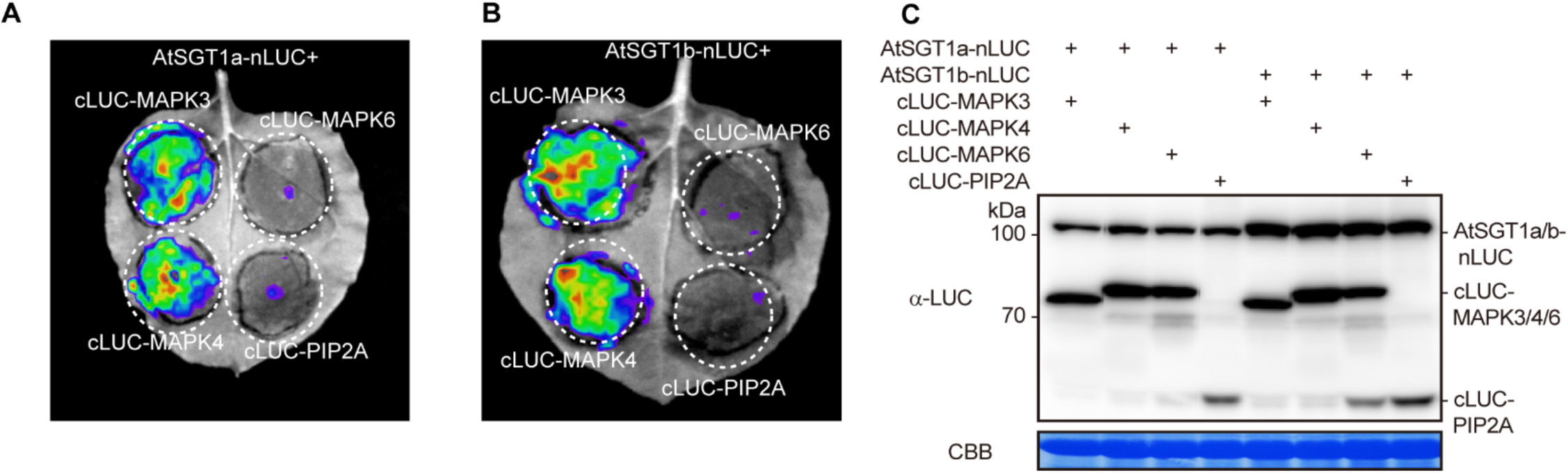
AtSGT1s associate with AtMAPKs. (A, B) AtSGT1a/b associate with AtMAPK3/4, but not with MAPK6, in Split-LUC assay. Agrobacterium combinations with different constructs were infiltrated in *N. benthamiana* leaves and luciferase activities were examined with CCD imaging machine. The MAPK-PIP2A combination was used as negative control. (C) Protein accumulation in (A) and (B).

**Supplemental Figure 10.**
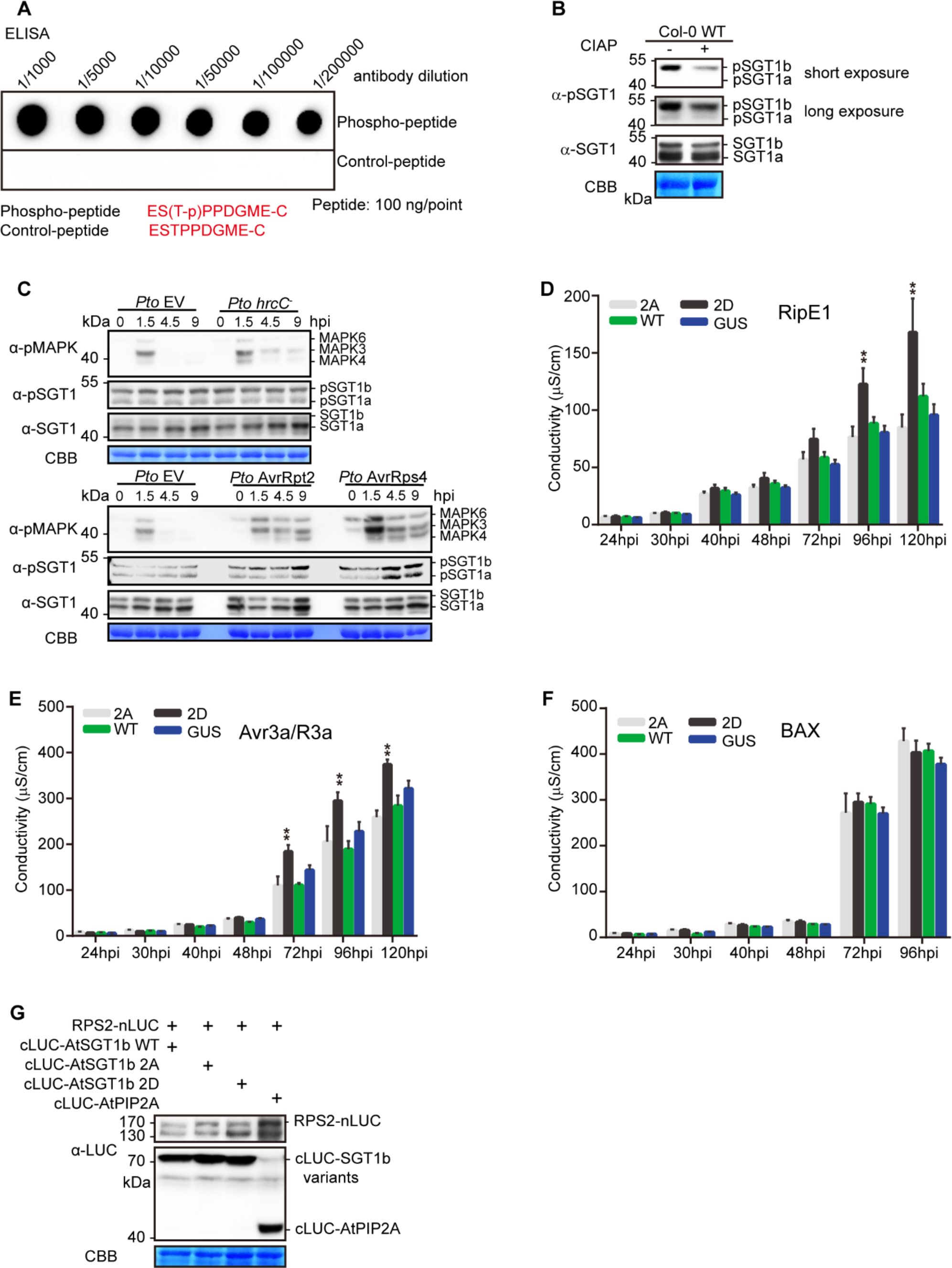
Phosphorylation of SGT1 T346 contributes to ETI robustness. (A) Determination of the specificity of the pSGT1 antibody using an *in vitro* ELISA assay. (B) Determination of the specificity of pSGT1 antibody in Arabidopsis using CIAP treatment. Two 12-d-old Col-0 WT plants were used for protein extract and CIAP enzyme treatment (37 °C, 60min) and the treated protein sample was subjected to western blot with the indicated antibodies. (C) ETI-triggered MAPK activation correlates with increased AtSGT1 phosphorylation at T346. Four-week-old Col-0 plants were infiltrated with *Pto* EV, *Pto hrcC^−^*, *Pto* AvrRpt2, or *Pto* AvrRps4 and samples were taken at the indicated time points. This experiment was performed twice with similar results. (D-F) A phospho-mimic mutation in AtSGT1b T346 (T346D) promotes cell death triggered by RipE1, Avr3a/R3a, but not BAX overexpression. Agrobacterium expressing AtSGT1b variants or the GUS-FLAG control (OD600=0.5) were infiltrated into *N. benthamiana* leaves 1 day before infiltration with Agrobacterium expressing RipE1, Avr3a/R3a, or BAX (OD600=0.15). Leaf discs were taken 21 hpi for conductivity measurements at the indicated time points. The time points in the x-axis are indicated as hpi with Agrobacterium expressing RipE1, Avr3a/R3a, or BAX (mean ± SEM, n=3, ** p<0.01, *t*-test, 3 replicates). (G) Western blot shows protein accumulation in Figure 5D and 5E.

**Supplemental Figure 11.**
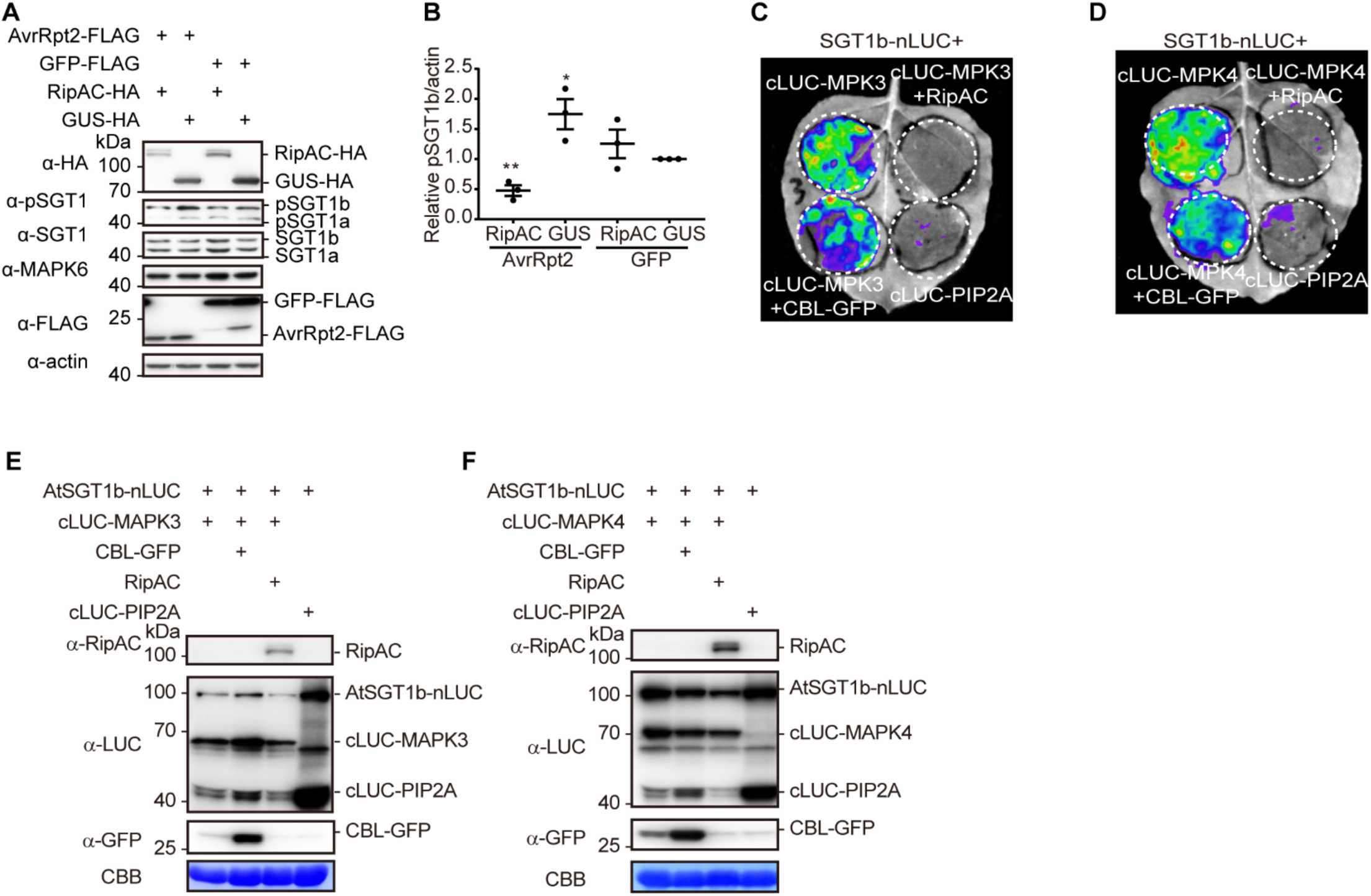
RipAC interferes with MAPK-SGT1 interaction to suppress SGT1 phosphorylation. (A) RipAC suppresses ETI-triggered SGT1 phosphorylation in Arabidopsis protoplasts. Western blots were performed as in Figure 4A using samples from protoplasts 10 hours after transfection with the indicated constructs. (B) Quantification of pSGT1b signal normalized to actin and relative to the control sample transfected with GUS and GFP in (A) (mean ± SEM of 3 independent biological replicates, * p<0.05, ** p<0.01, *t*-test). (C, D) Competitive Split-LUC assays showing that RipAC interferes with the interaction between MAPK3 (C) / MAPK4 (D) and AtSGT1b. Luciferase activity was determined with CCD camera. (E-F) Western blots shows protein accumulation in Figure 6E-F and S11E-F. These experiments were repeated at least 3 times with similar results.

**Supplemental Figure 12.**
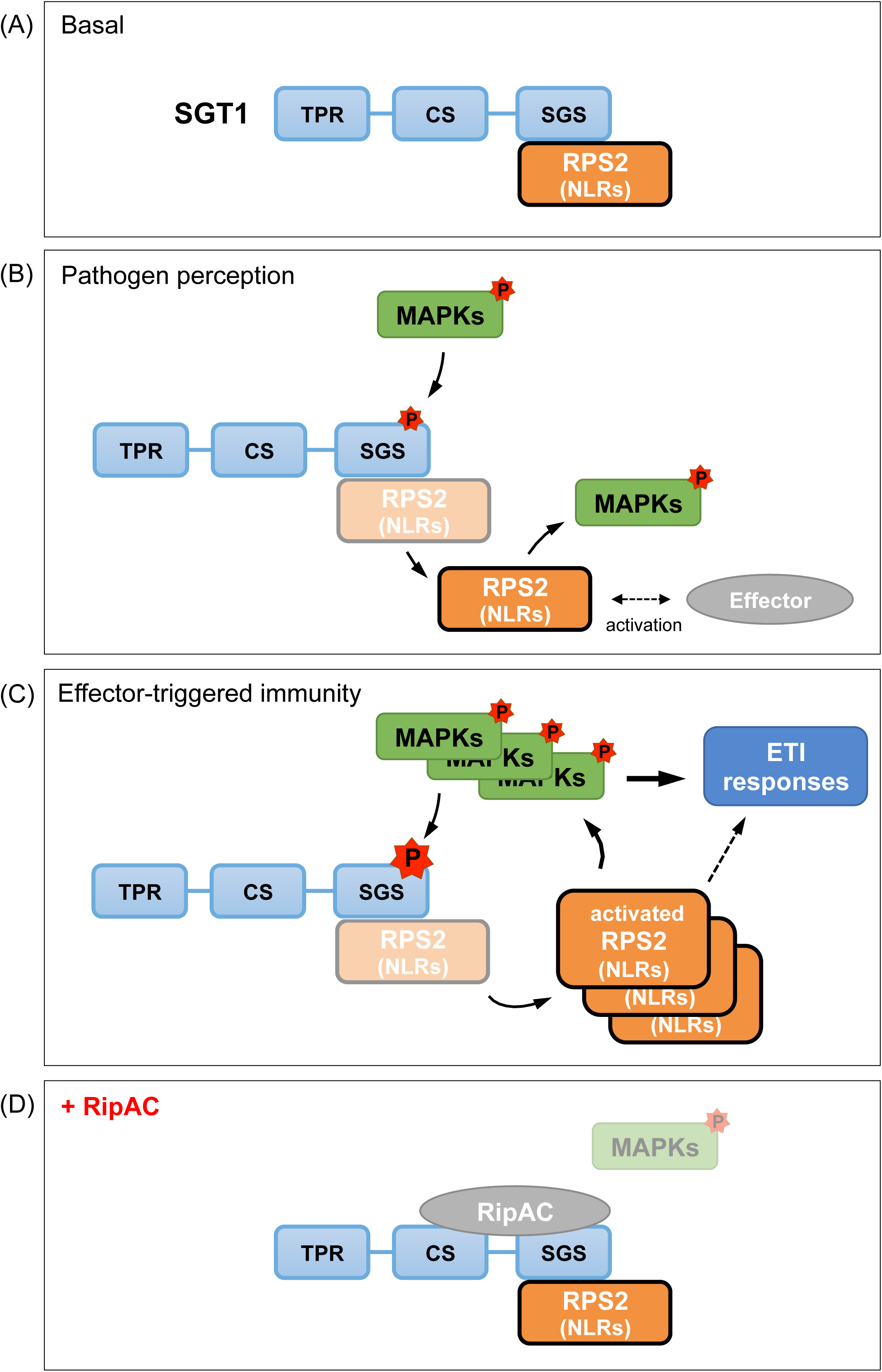
Simplified model to illustrate the activation of defense responses mediated by the phosphorylation of SGT1, and the impact of RipAC in this process. (A) In basal conditions, SGT1 (in blue, the diagram shows the different SGT1 domains) associates with NLRs (in our case, we studied RPS2), regulating NLR homeostasis and preventing NLR activation. (B) Pathogen perception leads to the phosphorylation and activation of MAPKs, likely derived from the activation of PRR-mediated signaling. MAPKs phosphorylate SGT1 SGS domain, relaxing the interaction between SGT1 and NLRs. Released NLRs can get activated (directly or indirectly) by pathogen effectors, leading to additional and sustained MAPK activation. (C) MAPK activation is known to contribute to the activation of ETI responses, while it mediates a feedback loop that enhances SGT1 phosphorylation. This, together with the transcriptional activation of *NLR* genes (triggered upon pathogen perception) leads to an enhanced pool of released activated NLRs. The activated NLR pool contributes to the sustained MAPK activation and, most likely, to the activation of ETI responses through other mechanisms. (D) RipAC interacts with the CS+SGS domains of SGT1. RipAC inhibits the MAPK-mediated phosphorylation of the SGT1 SGS domain. SGT1 phosphorylation at the SGT1 SGS domain is required for the release of NLRs.

**Supplemental Figure 13.**
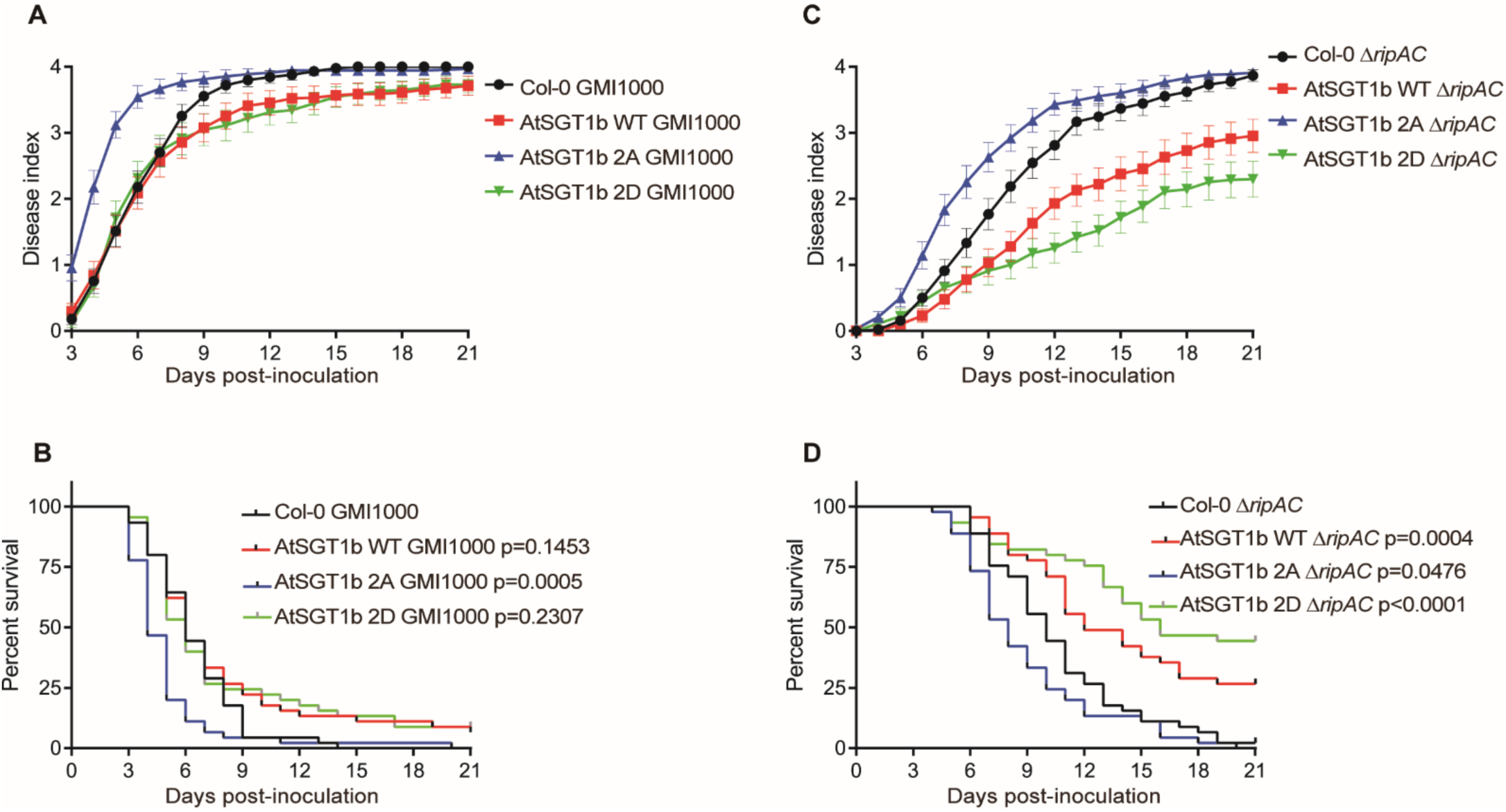
MAPK-mediated phosphorylation of AtSGT1b contributes to *Ralstonia solanacearum* resistance in Arabidopsis. (A, C) Soil-drenching inoculation assays in T3 generation AtSGT1b variants transgenic Arabidopsis were performed with GMI1000 WT (A) or Δ*ripAC* mutant (C) strains. The results are represented as disease progression, showing the average wilting symptoms in a scale from 0 to 4. Each data point represents the mean disease index for three independent experiments (n = 45 for each combination of bacterial strain and plant genotype). Each vertical bar represents the SEM from three independent experiments. (B, D) Survival analyses of Arabidopsis in (A, C). Statistical analysis were performed with Log-rank (Mantel-Cox) test (n = 45 for each combination of bacterial strain and plant genotype). The calculated P values relative to the Col-0 control are presented in the graphs.

