## Supplemental Materials for "A bacterial effector protein prevents MAPK-mediated phosphorylation of SGT1 to suppress plant immunity"

**This PDF file includes:**

Tables S1 to S2

SI References

Figures S1 to S13

**SI Appendix, Table S1. Resources used in this article**

| REAGENT or RESOURCE | SOURCE | IDENTIFIER |
| --- | --- | --- |
| Antibodies | | |
| Mouse monoclonal anti-GFP | Abiocode | Cat# M0802-3a |
| Mouse monoclonal anti-FLAG | Abmart | Cat# M20008 |
| Rabbit polyclonal anti-FLAG | Sigma | Cat# F7425 |
| Rabbit polyclonal anti-Luciferase | Sigma | Cat# L0159 |
| Phospho-p44/42 MAPK (Erk1/2) (Thr202/Tyr204)(20G11) Rabbit mAb antibody | Cell Signaling | Cat# 4370 |
| Mouse monoclonal anti-HA | Roche | Cat# 12CA5 |
| Rabbit polyclonal anti-MAPK6 | Agrisera | Cat# AS12 2633 |
| Rabbit polyclonal anti-actin | Agrisera | Cat# AS13 2640 |
| Rabbit polyclonal anti-AtSGT1a | (Yu et al., 2019) | N/A |
| Rabbit polyclonal anti-RipAC | This work | N/A |
| Rabbit polyclonal anti-SGT1 pT346 | This work | N/A |
| Rabbit polyclonal anti-H^+^-ATPase | Agrisera | Cat# AS07 260 |
| anti-Mouse IgG-Peroxidase | Sigma | Cat# A2554 |
| anti-Rabbit IgG-Peroxidase | Sigma | Cat# A0545 |
| Bacterial and Virus Strains | | |
| *Escherichia coli* DH5a | Transgen | CD501-3 |
| *Agrobacterium tumefaciens* GV3101 | Weidi Bio | AC1001 |
| *Ralstonia solanacearum* GMI1000 | (Salanoubat et al., 2002) | N/A |
| *Ralstonia solanacearum* GMI1000 *ΔripAC* | This work | N/A |
| *Ralstonia solanacearum* GMI1000 *ripAC^+^* | This work | N/A |
| *Pseudomonas syringae* pv. *tomato* (Pto)  DC3000 EV | (Macho et al., 2010) | N/A |
| *Pseudomonas syringae* pv tomato (Pto)  DC3000 *hrcC^-^* | (Ronald et al., 1992) |  |
| *Pseudomonas syringae* pv. *tomato* (Pto)  DC3000 AvrRpm1 | (Macho et al., 2010) | N/A |
| *Pseudomonas syringae* pv. *tomato* (Pto)  DC3000 AvrRpt2 | (Macho et al., 2010) | N/A |
| *Pseudomonas syringae* pv. *tomato* (Pto)  DC3000 AvrRps4 | (Macho et al., 2010) | N/A |
| Chemicals, Peptides, and Recombinant Proteins | | |
| Protease Inhibitor Cocktail for plant cell and tissue extracts, DMSO solution | Sigma | P9599 |
| GFP-Trap_A | Chromotek | Cat# gta-100 |
| ANTI-FLAG M2 Affinity Gel | Sigma-Aldrich | Cat# A2220 |
| XenoLight D-Luciferin | PerkinElmer | Cat# 122799 |
| Critical Commercial Assays |  |  |
| pENTR/D-TOPO Cloning Kit | Invitrogen | Cat# K240020SP |
| Gateway LR Clonase II Enzyme Mix | Invitrogen | Cat# 11791100 |
| Calf Intestinal Alkaline Phosphatase | New England Lab | Cat# NEB#M0290 |
| Experimental Models: Organisms/Strains | | |
| Arabidopsis: 35S:RipAC-GFP (AC#3 and AC #31) | This work | N/A |
| Arabidopsis: GVG:AvrRpt2 | (McNellis et al., 1998) | N/A |
| Arabidopsis: GVG:AvrRpt2/Col-0 | This work | N/A |
| Arabidopsis: GVG:AvrRpt2/AC #3 | This work | N/A |
| Arabidopsis: 35S:AtSGT1b WT | This work | N/A |
| Arabidopsis: 35S:AtSGT1b T346A (2A) | This work | N/A |
| Arabidopsis: 35S:AtSGT1b T346D (2D) | This work | N/A |
| *Solanum lycopersicum* cv. Moneymaker | N/A | N/A |
| Oligonucleotides | | |
| Primers see Table S2 | Ruidi Biotech | Custom order |
| Recombinant DNA | | |
| pENTR/D-TOPO | ThermoFisher | Cat# K240020 |
| pGWB502 | (Nakagawa et al., 2007) | N/A |
| pGWB505 | (Nakagawa et al., 2007) | N/A |
| pGWB511 | (Nakagawa et al., 2007) | N/A |
| pGWB554 | (Nakagawa et al., 2007) | N/A |
| pGTQL1211YN | (Lu et al., 2010) | Addgene # 61704 |
| pGTQL1221YC | (Lu et al., 2010) | Addgene # 61705 |
| pGWB-nLUC | (Wang et al., 2019) | N/A |
| pGWB-cLUC | This work | N/A |
| pEASYBLUNT-LB-Gm-RB | This work | N/A |
| pRCT-pRipAC-RipAC | This work | N/A |
| pGWB502-RipAC (no tag) | This work | N/A |
| pGWB505-RipAC-GFP | This work | N/A |
| pGWB-RipAC-nLUC | This work | N/A |
| pGWB-cLUC-RipAC | This work | N/A |
| pGWB502-CBL-GFP (no tag) | This work | N/A |
| 35S:RPS2-HA | (Li et al., 2014b) | N/A |
| 35S:R3a | (Liu et al., 2011) | N/A |
| 35S:Avr3a | (Liu et al., 2011) | N/A |
| 35S:Bax | (Liu et al., 2011) | N/A |
| 35S:INF1 | (Liu et al., 2011) | N/A |
| pGWB511-AtSGT1a-FLAG | This work | N/A |
| pGWB511-AtSGT1b-FLAG | This work | N/A |
| pGWB511-NbSGT1-FLAG | This work | N/A |
| pGWB511-SlSGT1b-FLAG | This work | N/A |
| pGWB554-AtSGT1a-RFP | This work | N/A |
| pGWB554-AtSGT1b-RFP | This work | N/A |
| pGWB554-NbSGT1-RFP | This work | N/A |
| pGWB-AtSGT1a-nLUC | This work | N/A |
| pGWB-AtSGT1b-nLUC | This work | N/A |
| pGWB-cLUC-AtSGT1a | This work | N/A |
| pGWB-cLUC-AtSGT1b | This work | N/A |
| pGWB-cLUC-NbSGT1 | This work | N/A |
| pGWB-cLUC-AtPIP2A | This work | N/A |
| pCAMBIA-AtFLS2-nLUC | (Li et al., 2014a) | N/A |
| pGWB512-FLAG-AtMAPK3 | This work | N/A |
| pGWB512-FLAG-AtMAPK4 | This work | N/A |
| pGWB512-FLAG-AtMAPK6 | This work | N/A |
| pGWB-cLUC-AtMAPK3 | This work | N/A |
| pGWB-cLUC-AtMAPK4 | This work | N/A |
| pGWB-cLUC-AtMAPK6 | This work | N/A |
| pGWB511-GUS-FLAG | (Yu et al., 2019) | N/A |
| pGTQL1211-RipAC-nYFP | This work | N/A |
| pGTQL1211-RipAC-cYFP | This work | N/A |
| pGTQL1211-AtPIP2A-nYFP | This work | N/A |
| pGTQL1211-AtPIP2A-cYFP | This work | N/A |
| pGTQL1211-AtSGT1a-cYFP | This work | N/A |
| pGTQL1211-AtSGT1b-cYFP | This work | N/A |
| pGTQL1211-NbSGT1-cYFP | This work | N/A |
| pGWB511-AtSGT1b TPR-FLAG | This work | N/A |
| pGWB511-AtSGT1b TPR+CS-FLAG | This work | N/A |
| pGWB511-AtSGT1b CS-FLAG | This work | N/A |
| pGWB511-AtSGT1b CS+SGS-FLAG | This work | N/A |
| pGWB511-AtSGT1b SGS-FLAG | This work | N/A |
| pCAMBIA-RPS2-nLUC | This work | N/A |
| pGWB502-AtSGT1b S271A (no tag) | This work | N/A |
| pGWB502-AtSGT1b S271D (no tag) | This work | N/A |
| pGWB502-AtSGT1b T346A (no tag) | This work | N/A |
| pGWB502-AtSGT1b T346D (no tag) | This work | N/A |
| pGWB502-AtSGT1b S271AT346A (no tag) | This work | N/A |
| pGWB502-AtSGT1b S271D T346D (no tag) | This work | N/A |
| pGWB-cLUC-AtSGT1b S271A | This work | N/A |
| pGWB-cLUC-AtSGT1b S271D | This work | N/A |
| pGWB-cLUC-AtSGT1b T346A | This work | N/A |
| pGWB-cLUC-AtSGT1b T346D | This work | N/A |
| pGWB-cLUC-AtSGT1b S271AT346A | This work | N/A |
| pGWB-cLUC-AtSGT1b S271D T346D | This work | N/A |
| pGWB-505-GFP | This work | N/A |
| pGWB-505-AtSGT1b-GFP | This work | N/A |
| pGWB-505-AtSGT1b S271AT346A-GFP | This work | N/A |
| pGWB-505-AtSGT1b S271D T346D-GFP | This work | N/A |
| pGWB505-AtMKK7DD-GFP | This work | N/A |
| pGWB505-RipE1 | (Sang et al., 2019) | N/A |
| pGWB-cLUC-AtPIP2A | This work | N/A |
| pHBT-AvrRpt2-FLAG | This work | N/A |
| pHBT-GFP-FLAG | This work | N/A |
| pXCSG-RipAC-HA-Strep | This work | N/A |
| pXCSG-GUS-HA-Strep | This work | N/A |
| Software and Algorithms |  |  |
| Prism 7 | GraphPad Software | https://www.graphpad.com/scientific-software/prism/ |
| Scaffold 4.0 | Proteome Software | http://www.proteomesoftware.com/products/scaffold/ |
| Adobe Illustrator CS6 (64Bit) | Adobe Illustrator Software | https://www.adobe.com/products/illustrator.html |
| ImageJ | NIH ImageJ | https://imagej.nih.gov/ij/ |

**SI Appendix, Table S2. Primers used in this study.**

| Name | Sequence 5’ to 3’ | Purpose |
| --- | --- | --- |
| RipAC-LB-F | CGATTTCACGCCGGCGAAGAC | Clone *RipAC* left board |
| RipAC-LB-R | CCGGATCAAGAATTCGAGCGTCAGGCTAAGTCGCG |  |
| RipAC-RB-F | CCTGACGCTCGAATTCTTGATCCGGTGCCGCATCCC | Clone *RipAC* right board |
| RipAC-RB-R | CGACCAGGCGCGCATCACC |  |
| pRipAC-F | CCCCCTAGGTCACTTATTGGAAACTCCCGTC | Clone *RipABC* operon promoter |
| pRipAC-R | GGGGTACCCAGGCTCCGGCGGGCCGGAGCGC |  |
| RipAC-F | CACCATGCCTATCCTTCCACGCCTATTCCATCGA | Clone *RipAC* CDS |
| RipAC-R | ACGCTGCCTCGACGGACTTGCCGCGGGCGT |  |
| AtSGT1a-F | CACCATGGCGAAGGAGCTTGCT | Clone *AtSGT1a* CDS |
| AtSGT1a-R | GATCTCCCATTTCTTGAGCTCCAT |  |
| AtSGT1b-F | CACCATGGCCAAGGAATTAGCAGAGAAAGCTAAAGAAGCT | Clone *AtSGT1b* CDS |
| AtSGT1b-R | ATACTCCCACTTCTTGAGCTCCATGCCATCTG |  |
| NbSGT1-F | CACCATGGCGTCCGATCTGGAGATTAGGGC | Clone *NbSGT1* CDS |
| NbSGT1-R | GATTTCCCATTTCTTCAGCTCCATGCC |  |
| SlSGT1b-F | CACCATGGCGTCCGATCTGGAGACTAGGGCTAAA | Clone *SlSGT1b* CDS |
| SlSGT1b -R | GATCTCCCATTTCTTCAGCTCCATGCC |  |
| AtPIP2A-F | CACCATGGCAAAGGATGTGGAAGCCGTTCCCG | Clone *AtPIP2A* CDS |
| AtPIP2A-R | GACGTTGGCAGCACTTCTGAATGATCCG |  |
| CBL-GFP-F | CACCATGGGCTGCTTCCACTCAAAGGCAGCAAAAGAATTTATGGTGAGCAAGGGCGAGGAGCTGTTCA | Clone *CBL-GFP* CDS |
| GFP-R | CTTGTACAGCTCGTCCATGCCGTGAGTG |  |
| AvrRpt2-F | ATGAAAATTGCTCCAGTTGCCAT | RT-PCR |
| AvrRpt2-R | GTAGAGCATTGCGTGTGGAAC | RT-PCR |
| Atactin2-F | TGCTGGACGTGACCTTACTG | RT-PCR |
| Atactin2-R | TTCTCGATGGAAGAGCTGGT | RT-PCR |
| AtSGT1b S271A-F | ttttgctggcttagaagctgggtacactggtctct | Generate *AtSGT1b* mutant (S271A) |
| AtSGT1b S271A-R | agagaccagtgtacccagcttctaagccagcaaaa |  |
| AtSGT1b S271D-F | gtcttttgctggcttagaatctgggtacactggtctctgc | Generate *AtSGT1b* mutant (S271D) |
| AtSGT1b S271D-R | gcagagaccagtgtacccagattctaagccagcaaaagac |  |
| AtSGT1b T346A-F | catgccatctggtggagcgctctccactttcttag | Generate *AtSGT1b* mutant (T346A) |
| AtSGT1b T346A-R | ctaagaaagtggagagcgctccaccagatggcatg |  |
| AtSGT1b T346D-F | ctccatgccatctggtggatcgctctccactttcttagtc | Generate *AtSGT1b* mutant (T346D) |
| AtSGT1b T346D-R | gactaagaaagtggagagcgatccaccagatggcatggag |  |
| NbSGT1 TPR-R | GGCAGCAGATCCTTGATAGGACAG | Generate *NbSGT1* TPR truncation |
| NbSGT1 CS-F | CACCATGTCTGAGTCTTTGGGCAATGTTGCTG | Generate *NbSGT1* CS truncation |
| NbSGT1 CS-R | AGACTCTCTCGTATATTCGAGAGATG |  |
| NbSGT1 SGS-F | CACCATGTATACGAGAGAGTCTGCTGTAGTGC | Generate *NbSGT1* SGS truncation |
| AtMAPK4-F | CACCATGTCGGCGGAGAGTTGTTTCGGAAGCTCG | Clone *AtMAPK4* CDS |
| AtMAPK4-R | CACTGAGTCTTGAGGATTGAACTTGACT |  |
| ATMKK7-F | caccATGGCTCTTGTTCGTAAACGC | Clone *AtMKK7* CDS |
| ATMKK7-R | AAGACTTTCACGGAGAAAAGGG |  |
| MKK7S193D-F | gacgtaggaattgcagtaatctaaatctcgggtaatgattttgctcactcc | Generate *AtMKK7* mutant (S193D) |
| MKK7S193D-R | ggagtgagcaaaatcattacccgagatttagattactgcaattcctacgtc |  |
| MKK7S199D-F | tacccgatcgttagattactgcaatgactacgtcggcacttg | Generate *AtMKK7* mutant (S199D) |
| MKK7S199D-R | caagtgccgacgtagtcattgcagtaatctaacgatcgggta |  |
| pHBT-AvrRpt2- FLAG-F | CTTGCTCCGTGGATCCTCTAGAATGAAAATTGCTCCAGTTGCC | Clone AvrRpt2 into pHBT-35S-FLAG |
| pHBT-AvrRpt2- FLAG-R | TGTAGTCAGAAGGCCTGGTACCGCGGTAGAGCATTGCGTGTG |  |
| pHBT-GFP- FLAG-F | CTTGCTCCGTGGATCCTCTAGAATGGTGAGCAAGGGCGA | Clone GFP into pHBT-35S-FLAG |
| pHBT-GFP- FLAG-R | TGTAGTCAGAAGGCCTGGTACCCTTGTACAGCTCGTCCATG |  |
| RPS2-nLUC-F | gagaacacgggggacgagctcATGGATTTCATCTCATCTCTTATCG | Clone RPS CDS into pCAMBIA-nLUC |
| RPS2-nLUC-R | cgcgtacgagatctggtcgacATTTGGAACAAAGCGCGGT |  |

**Methods references**

Li, L., Li, M., Yu, L., Zhou, Z., Liang, X., Liu, Z., Cai, G., Gao, L., Zhang, X., Wang, Y., et al. (2014a). The FLS2-associated kinase BIK1 directly phosphorylates the NADPH oxidase RbohD to control plant immunity. Cell host & microbe 15:329-338.

Li, M., Ma, X., Chiang, Y.-H., Yadeta, K.A., Ding, P., Dong, L., Zhao, Y., Li, X., Yu, Y., Zhang, L., et al. (2014b). Proline isomerization of the immune receptor-interacting protein RIN4 by a cyclophilin inhibits effector-triggered immunity in Arabidopsis. Cell host & microbe 16:473-483.

Liu, T., Ye, W., Ru, Y., Yang, X., Gu, B., Tao, K., Lu, S., Dong, S., Zheng, X., Shan, W., et al. (2011). Two host cytoplasmic effectors are required for pathogenesis of *Phytophthora sojae* by suppression of host defenses. Plant Physiology 155:490-501.

Lu, Q., Tang, X., Tian, G., Wang, F., Liu, K., Nguyen, V., Kohalmi, S.E., Keller, W.A., Tsang, E.W.T., Harada, J.J., et al. (2010). Arabidopsis homolog of the yeast TREX-2 mRNA export complex: components and anchoring nucleoporin. The Plant Journal 61:259-270.

Macho, A.P., Guevara, C.M., Tornero, P., Ruiz-Albert, J., and Beuzón, C.R. (2010). The *Pseudomonas syringae* effector protein HopZ1a suppresses effector-triggered immunity. New Phytologist 187:1018-1033.

McNellis, T.W., Mudgett, M.B., Li, K., Aoyama, T., Horvath, D., Chua, N.-H., and Staskawicz, B.J. (1998). Glucocorticoid-inducible expression of a bacterial avirulence gene in transgenic Arabidopsis induces hypersensitive cell death. The Plant Journal 14:247-257.

Nakagawa, T., Suzuki, T., Murata, S., Nakamura, S., Hino, T., Maeo, K., Tabata, R., Kawai, T., Tanaka, K., Niwa, Y., et al. (2007). Improved gateway binary vectors: high-performance vectors for creation of fusion constructs in transgenic analysis of plants. Bioscience, Biotechnology, and Biochemistry 71:2095-2100.

Ronald, P.C., Salmeron, J.M., Carland, F.M., and Staskawicz, B.J. (1992). The cloned avirulence gene *avrPto* induces disease resistance in tomato cultivars containing the *Pto* resistance gene. Journal of bacteriology 174:1604-1611.

Salanoubat, M., Genin, S., Artiguenave, F., Gouzy, J., Mangenot, S., Arlat, M., Billault, A., Brottier, P., Camus, J.C., Cattolico, L., et al. (2002). Genome sequence of the plant pathogen *Ralstonia solanacearum*. Nature 415:497-502.

Sang, Y., Yu, W., Zhuang, H., Wei, Y., Derevnina, L., Luo, J., Kamoun, S., and Macho, A.P. (2019). Intra-strain elicitation and suppression of plant immunity by *Ralstonia solanacearum* type-III effectors in *Nicotiana benthamiana*. bioRxiv:780890.

Wang, Y., Li, Y., Rosas-Diaz, T., Caceres-Moreno, C., Lozano-Durán, R., and Macho, A.P. (2019). The IMMUNE-ASSOCIATED NUCLEOTIDE-BINDING 9 protein is a regulator of basal immunity in *Arabidopsis thaliana*. Molecular Plant-Microbe Interactions 32:65-75.

Yu, G., Xian, L., Sang, Y., and Macho, A.P. (2019). Cautionary notes on the use of Agrobacterium-mediated transient gene expression upon SGT1 silencing in *Nicotiana benthamiana*. New Phytologist 222:14-17.
